# Th17 cells require the DNA repair sensor XPC to control oxidative DNA damage

**DOI:** 10.1101/2024.09.13.612834

**Authors:** Jefferson Antônio Leite, Natalia Notaberardino Bos, Luísa Menezes, Eloísa Martins, Giovana da Silva Leandro, Patrick da Silva, Hedden Ranfley, Samuel Oliveira, Camila Pontes Ferreira, José Arimatéia de Oliveira Nery Neto, Victor Yuji, Marcela Cipelli, Beatriz Leocata, Talita Gonçalves, Anthony Gabry da Silveira, Sabrina Baroni, Howard L. Weiner, Thomas Hofmann, Stefanie Scheu, Leandro Machado Colli, Ari Waisman, Sandra Muxel, Carlos Frederico Martins Menck, Niels Olsen Saraiva Câmara

## Abstract

Th17 cells are critical for mucosal immunity, producing IL-17A, IL-17F, and IL-22, but dysregulated Th17 responses are implicated in autoimmune diseases. Despite their susceptibility to oxidative stress in certain conditions, Th17 cells exhibit reduced oxidative DNA damage and cell death compared to other T helper subsets. However, the mechanisms that protect Th17 cells from oxidative stress are poorly understood. Here, we identify Xeroderma Pigmentosum Complementation Group C (XPC) as a key regulator of DNA repair and genomic stability in Th17 cells. In XPC-deficient mice, we demonstrate that the absence of XPC impairs Th17 differentiation, as evidenced by reduced expression of key differentiation markers, including Rorc and Il17a, along with decreased IL-17A production. This deficiency leads to increased oxidative stress, DNA damage, and a metabolic shift from glycolysis to oxidative phosphorylation. Moreover, the transcription factor BATF directly regulates XPC expression, linking the BATF-XPC axis to the maintenance of Th17 cell function. Importantly, we find that restoring antioxidant capacity with N-Acetylcysteine (NAC) rescues IL-17A production and reduces DNA damage in XPC-deficient Th17 cells. Mechanistically, we find that XPC interacts with OGG1, a DNA glycosylase involved in the repair of oxidative DNA damage, highlighting XPC’s role in maintaining genomic integrity during Th17 cell differentiation. Our findings reveal a previously unrecognized role for XPC in protecting Th17 cells from oxidative stress, ensuring their proper differentiation and function, with potential implications for targeting DNA repair pathways in autoimmune and inflammatory diseases.

## Introduction

T helper 17 (Th17) cells are a subset of T helper cells characterized by the production of IL-17A, IL-17F and IL-22 cytokines. They are important for a protective immunity against bacterial and fungi infections at mucosal surfaces. However, dysregulated Th17 cell responses are involved in the pathogenesis of several inflammatory or autoimmune diseases, such as inflammatory bowel disease, psoriasis and multiple sclerosis ^1^. The development of Th17 cells relies on the presence of cytokines such as transforming growth factor-β (TGF-β) and IL-6. TGF-β triggers the activation of Smad2/3 transcription factors, while IL-6 signals facilitate STAT3 phosphorylation. The concerted action of Smad2/3 and STAT3, along with other transcription factors activated by TCR signaling, promotes the expression of RORγt and facilitates Th17 cell differentiation. IL-21 and IL-23 can also transmit signals through STAT3, playing a crucial role in the differentiation of both murine and human Th17 cells ^2–6^.

Endogenous factors threaten genome stability and alter T cell differentiation, which impact on autoimmunity and cancer development ^7–9^. Reactive oxygen species (ROS) have been considered harmful products from metabolism, which damage cellular proteins, lipids, and nucleic acids^10^. Activated T cells can induce respiratory bursts both through direct contact with phagocytes and via the release of cytokines^11^. TCR ligation triggers the activation of NOX-2, leading to the production of hydrogen peroxide (H_2_O_2_) that activates GATA-3 and STAT6, while inhibiting STAT3, therefore promoting Th2 and inhibiting Th17 cells differentiation^12^. In Th17 cells the absence of calcium signaling by STIM1s increases ROS production which results in DNA damage and cell death ^13^. On the other hand, ROS scavenger N-acetylcysteine boosts Th17 cell differentiation^14^. Despite being susceptible to increased amounts of ROS in the absence of calcium signaling, WT Th17 cells display less accumulation of ROS, DNA damage and cell death when compared to Th0 and Th1 cells ^15^, which suggest that Th17 cells could have mechanisms to protect against oxidative DNA damage.

The repair of oxidative-induced DNA damage occurs through sophisticated pathways that include the base excision repair (BER) and the nucleotide excision repair (NER). DNA glycosylases, such as OGG1 and NEIL family members, recognize and excise oxidized or damaged bases. Following this, Apurinic/apyrimidinic (AP) endonucleases, like APE1, cleave the DNA strand at the apurinic or apyrimidinic sites, creating a nick. DNA polymerases, notably DNA polymerase β, fill in the gap with newly synthesized DNA, and ligases, including DNA ligase III, seal the nick by forming phosphodiester bonds ^16^. The NER pathway is initiated by the Xeroderma Pigmentosum Complementation Group C (XPC) protein, which is crucial for the recognition of bulky DNA lesions, such as those caused by UV radiation. While XPC plays a key role in damage recognition, it indirectly contributes to the repair of oxidative DNA damage by facilitating the removal of structurally significant lesions, thereby ensuring the initiation of a precise and efficient repair process ^17–20^. Interestingly, Th17 cells upregulate genes associated with NER DNA repair pathway in the colon during intestinal inflammation^22^, however, it is unknown whether Th17 cells use DNA repair as a mechanism to control DNA damage. Therefore, we hypothesize that Th17 cells use the NER DNA repair pathway to control oxidative DNA damage and genomic instability contributing to an efficient differentiation and function.

Here, we demonstrate that Th17 cells upregulate different genes from the NER DNA repair pathway, including XPC in a BATF-dependent manner, and accumulate less DNA damage when compared to Th0 cells. The lack of XPC is associated with compromised Th17 cell differentiation, increased ROS levels and oxidative-induced DNA damage as well as a metabolic reprogramming change, namely less glycolytic capacity and more oxidative phosphorylation. XPC interacts with OGG1 to control oxidative-induced DNA damage in Th17 cells. Thus, XPC regulates oxidative DNA repair by modulating key DNA repair molecules and maintaining antioxidant responses. Thus, XPC is vital for protecting Th17 cells from oxidative DNA damage, ensuring their proper function.

## Results

### XPC is required for Th17 cells differentiation

Initially, we investigated whether the deficiency of XPC affects the differentiation of CD4+ and CD8+ T cells under steady-state conditions. We collected draining lymph nodes and spleens from 6-8 week-old WT and XPC^-/-^ mice. Flow cytometry analysis revealed no significant difference in the frequency of CD4+ and CD8+ T cells between WT and XPC^-/-^ mice in the lymph nodes and spleens **(Figure 1A-B)**. Moreover, the frequency of naive (CD62L+CD44-) and memory (CD62L-CD44+) subsets of both CD4+ and CD8+ T cells was comparable between the two groups **(Figure 1C-D)**.

**Figure 1:**
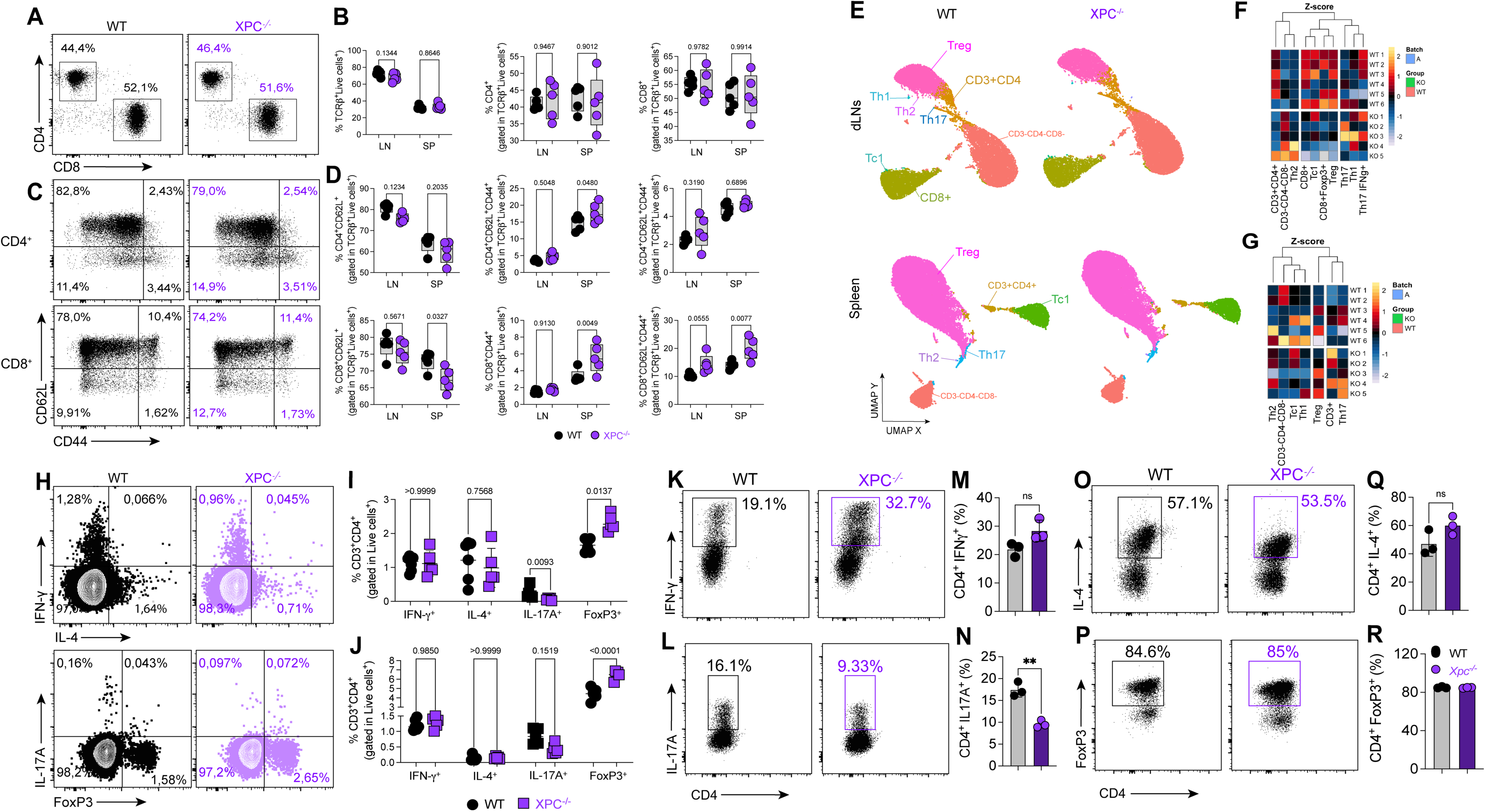
Flow cytometry and UMAP analysis of T cell subsets in WT and XPC-/- mice, and in vitro polarization of CD4+ T cells. **(A)** Flow cytometry analysis of CD4+ and CD8+ T cell populations from lymph nodes (LN) and spleens (SP) of WT and XPC-/- mice. Percentages represent the proportion of CD4+ and CD8+ cells within total T cells. **(B)** Quantification of the percentages of CD4+ and CD8+ T cells in LN and SP from WT and XPC-/- mice. **(C)** Flow cytometry plots showing CD62L and CD44 expression on CD4+ and CD8+ T cells from WT and XPC-/- mice, representing naïve (CD62L+CD44-) and memory (CD62L-CD44+) subsets. **(D)** Quantification of naïve and memory T cell subsets (CD62L+CD44- and CD62L-CD44+) in CD4+ and CD8+ T cells from LN and SP of WT and XPC-/- mice. **(E)** UMAP projections showing the clustering of T cell subsets from draining lymph nodes and spleens in WT and XPC-/- mice. Identified clusters include Th1, Th2, Th17, Tregs, and CD8+ T cells. **(F)** Heatmap displaying Z-scores for the expression of key transcription factors and cytokines in T cell subsets from LN of WT and XPC-/- mice. **(G)** Heatmap of Z-scores for transcription factors and cytokines in T cell subsets from spleens of WT and XPC-/- mice. **(H)** Flow cytometry plots showing intracellular staining for IFN-γ, IL-4, IL-17A, and FoxP3 in CD4+ T cells from LN of WT and XPC-/- mice. **(I)** Quantification of CD4+ T cells expressing IFN-γ, IL-4, IL-17A, and FoxP3 from LN of WT and XPC-/- mice. **(J)** Quantification of CD4+ T cells expressing IFN-γ, IL-4, IL-17A, and FoxP3 from SP of WT and XPC-/- mice. **(K)** Flow cytometry analysis of IFN-γ production in CD4+ T cells from in vitro Th1-polarized cultures of naïve CD4+ T cells isolated from WT and XPC-/- mice, stimulated with anti-CD3/CD28 under Th1-polarizing conditions. **(L)** Flow cytometry analysis of IL-17A production in CD4+ T cells from in vitro Th17-polarized cultures of naïve CD4+ T cells isolated from WT and XPC-/- mice, stimulated with anti-CD3/CD28 under Th17-polarizing conditions. **(M)** Bar graph showing the percentage of IFN-γ+ CD4+ T cells from in vitro Th1-polarized cultures of WT and XPC-/- mice. **(N)** Bar graph showing the percentage of IL-17A+ CD4+ T cells from in vitro Th17-polarized cultures of WT and XPC-/- mice. **(O)** Flow cytometry analysis of FoxP3 expression in CD4+ T cells from in vitro Treg-polarized cultures of naïve CD4+ T cells isolated from WT and XPC-/- mice, stimulated under Treg-polarizing conditions**. (P)** Flow cytometry analysis of FoxP3 expression in CD4+ T cells from in vitro Treg-polarized cultures of WT and XPC-/- mice. **(Q)** Bar graph showing the percentage of IL-4+ CD4+ T cells from in vitro Th2-polarized cultures of WT and XPC-/- mice. **(R)** Bar graph showing the percentage of FoxP3+ CD4+ T cells from in vitro Treg-polarized cultures of WT and XPC-/- mice. Data are representative of at least two independent experiments and are shown as mean ± SEM. *p < 0.05 determined T-test *p < 0.05

We next examined the differentiation of specific T cell subsets, including Th1, Th2, Th17, Treg, and Tc1 cells. UMAP plots confirmed that the overall distribution of these subsets was similar in both WT and XPC^-/-^ mice in the draining lymph nodes and spleen **(Figure 1E)**. Heatmaps comparing Z-scores of gene expression also confirmed no significant differences in the differentiation of these subsets **(Figure 1F-G)**. However, flow cytometry analysis showed specific differences in the frequencies of certain T cell subsets, particularly Th17 and Treg cells. In the lymph nodes, the percentage of IL-17A-producing CD4+ Th17 cells was significantly reduced in XPC**-/-** mice compared to WT controls **(Figure 1H, I), upper panels)**. This reduction in Th17 cells was specific to the lymph nodes, while no significant difference was observed in the spleen **(Figure 1H, J, lower panel)**. Interestingly, we observed a significant increase in the percentage of FoxP3+ regulatory T cells (Tregs) in both the lymph nodes and spleens of XPC^-/-^ mice compared to WT mice **(Figure 1H, I, J)**.

To further investigate whether XPC deficiency affects *in vitro* T helper cell differentiation, we isolated CD4^+^CD62L^high^ T cells from the lymph nodes and spleens of both WT and XPC^-/-^ mice and cultured them under Th1, Th2, Treg, and Th17 cell polarizing conditions for five days. The flow cytometry analysis showed that the deficiency of XPC did not affect the differentiation of Th1, Th2 and Treg cells **(Figure 1K, M, O, Q, P, R)**. However, the *in vitro* differentiation of Th17 cells was significantly impaired in the absence of XPC, as the frequency of IL-17A-producing cells was notably lower compared to WT mice **(Figure 1L, N)**. This reduction in Th17 differentiation was associated with diminished mRNA expression of *Rorc*, *Stat3*, and *Il17a*, as shown by qPCR analysis **(Supplementary Figures 1A-C)**.

In summary, while XPC deficiency does not impact the overall differentiation of CD4+ or CD8+ T cells, nor the development of Th1 or Th2 cells, it significantly impairs the generation of IL-17A-producing Th17 cells. Interestingly, XPC deficiency leads to an increase in FoxP3+ Treg cells in both the lymph nodes and spleen. These results suggest a critical role for XPC in the balance between Treg and Th17 populations under steady-state conditions.

### Th17 cells accumulate less DNA damage with enhanced XPC expression

Next, we investigated the role of XPC in the DNA damage response across different T cell subsets and explored potential regulatory mechanisms controlling its expression. We first measured the levels of γH2AX, a marker of DNA damage, across various T helper (Th) cell subsets and Treg cells using flow cytometry. As shown in Figure 2A, the naïve T cells displayed the lowest levels of γH2AX, indicating minimal DNA damage in these resting cells. In contrast, Th0 and Th2 cells showed significantly elevated γH2AX expression, reflecting higher levels of DNA damage or an active DNA repair response in these subsets. Interestingly, Th1, Th17, and Treg cells exhibited lower γH2AX levels compared to Th0 cells, though their expression was still higher than that of naïve T cells **(Figure 2A, B, C)**. This suggests that while Th1, Th17, and Treg cells engage DNA repair mechanisms, the extent of DNA damage (or repair activity) in these cells is less pronounced than in Th0 and Th2 cells.

**Figure 2:**
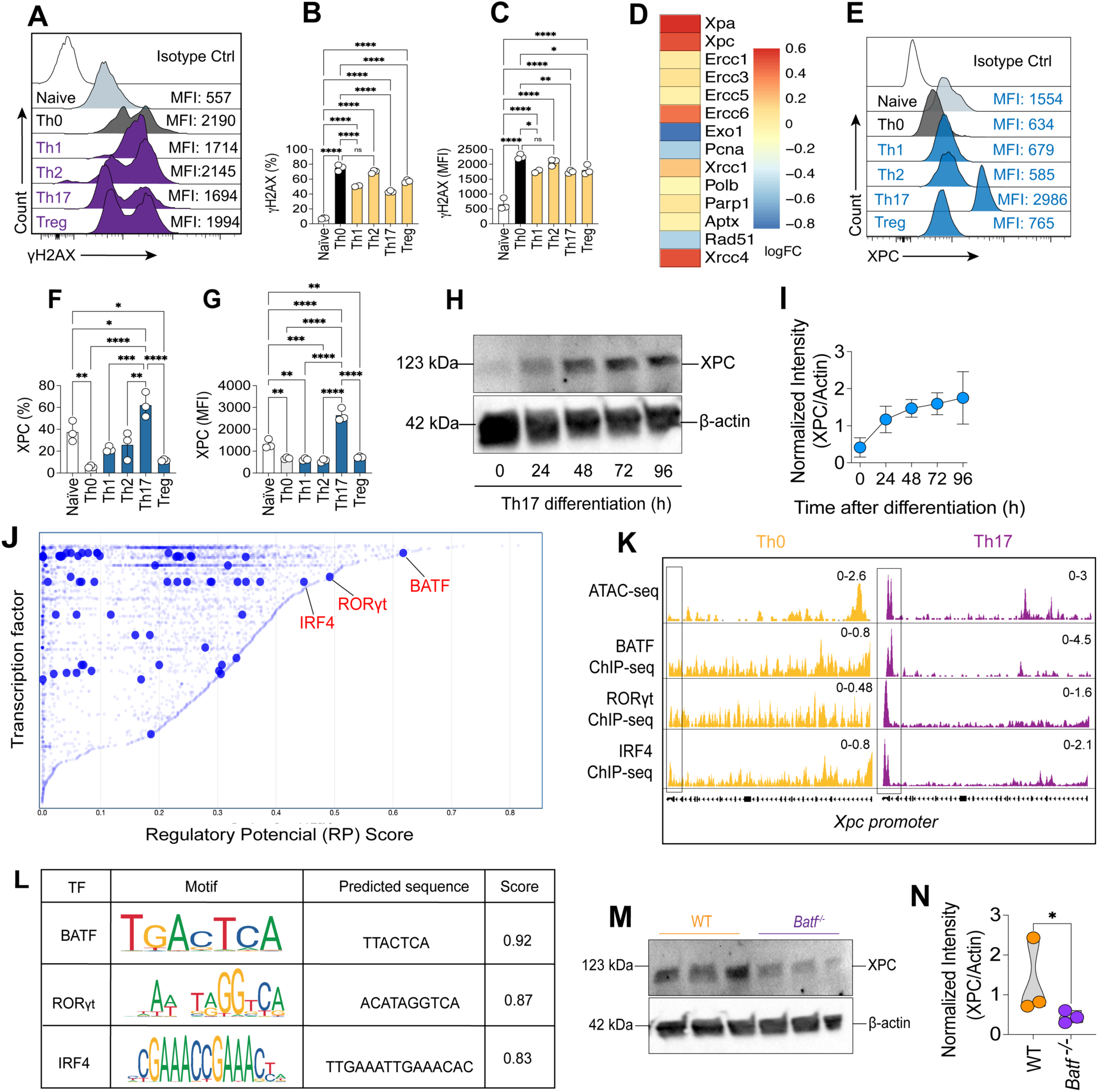
XPC expression and DNA damage response during in vitro T cell differentiation. **(A)** Flow cytometry histograms showing γH2AX expression in naïve, Th0, Th1, Th2, Th17, and Treg cells. Median fluorescence intensity (MFI) values are indicated for each subset. **(B)** Quantification of γH2AX expression (MFI) in naïve and polarized Th subsets (Th0, Th1, Th2, Th17, Treg). Data are shown as mean ± SD. Statistical significance was determined by one-way ANOVA. **(C)** Quantification of γH2AX expression (MFI) in CD4+ T cells isolated from WT and XPC-/- mice, under Th0, Th1, Th2, Th17, and Treg polarizing conditions. **(D)** Heatmap showing the logFC (fold change) of DNA repair-related genes in polarized Th subsets, including Xpa, Xpc, and others. **(E)** Flow cytometry histograms depicting XPC expression in naïve, Th0, Th1, Th2, Th17, and Treg cells. MFI values are indicated for each subset. **(F)** Percentage of XPC+ cells in naïve and polarized Th subsets, presented as mean ± SD. **(G)** Quantification of XPC expression (MFI) in naïve and polarized Th subsets. Data are shown as mean ± SD. **(H)** Western blot analysis of XPC expression during Th17 differentiation over a 96-hour time course. β-actin was used as a loading control. **(I)** Quantification of XPC protein levels during Th17 differentiation, normalized to β-actin. **(J)** Scatter plot showing the regulatory potential (RP) scores of transcription factors associated with Th17 differentiation. Key transcription factors such as BATF, RORγt, and IRF4 are highlighted. **(K)** ATAC-seq (Svensson MN, et al, 2019) and ChIP-seq tracks for BATF, RORγt, and IRF4 binding at the Xpc promoter in Th0 and Th17 conditions (Ciofani M, et al 2012). Analyses were performed using the Cistrome Project platform - Cistrome DB toolkit; **(L)** Predicted binding motifs for BATF, RORγt, and IRF4, along with their respective binding sequences and scores. **(M)** Western blot analysis of XPC expression in Th17 cells from WT and Batf-/- mice. β-actin was used as a loading control**. (N)** Quantification of XPC protein levels in WT and Batf-/- Th17 cells, normalized to β-actin. Data are representative of at least two independent experiments and are shown as mean ± SEM. *p < 0.05 determined by one-way ANOVA followed by Tukey’s post hoc test.

We then examined the expression of XPC and other DNA repair genes across Th17 cells compared to non-polarized Th0 cells. Heatmap analysis of log fold changes (logFC) showed that Th17 cells had increased expression of several DNA repair genes, including *Xpc*, *Xpa*, *Ercc1, Ercc3, Ercc5, Ercc6, Xcrcc1, Polb, Parp1, Aptx* and *Xcc4* when compared to Th0 cells **(Figure 2D)**. These findings suggest that DNA repair pathways, particularly nucleotide excision repair (NER) and other related mechanisms, are upregulated in Th17 cells. Flow cytometry confirmed these results, showing higher XPC frequency and MFI values in Th17 cells compared to naive T cells and other subsets **(Figure 2E, F, G)**.

We next sought to identify which transcription factors might regulate XPC expression in Th17 cells. To this end, we used regulatory potential (RP) scores, which rank transcription factors based on their likelihood of regulating specific genes. This analysis highlighted several key transcription factors, including BATF, RORγt, and IRF4, that exhibited high RP scores and are known to be critical for Th17 differentiation **(Figure 2J)**. Chromatin immunoprecipitation followed by sequencing (ChIP-seq) confirmed the binding of BATF, RORγt, and IRF4 to the *Xpc* promoter specifically in Th17 cells, with BATF showing particularly strong enrichment **(Figure 2K)**. This indicates that these transcription factors likely play a central role in driving *Xpc* expression during Th17 differentiation. Notably, the accessibility of the *Xpc* promoter (as indicated by ATAC-seq) was significantly higher in Th17 cells compared to non-polarized Th0 cells, underscoring the transcriptional activation of *Xpc* during Th17 lineage commitment. Motif analysis further identified BATF as a key regulator, with a high affinity binding site on the *Xpc* promoter. This was supported by the motif scores, which showed BATF as having the highest predicted binding sequence among the transcription factors analyzed, followed by RORγt and IRF4**(Figure 2L)**. These results suggest that BATF, in cooperation with RORγt and IRF4, directly regulates Xpc transcription in Th17 cells, potentially coordinating the DNA repair machinery needed during active cell differentiation.

To validate the role of BATF in controlling XPC expression, we examined XPC levels in BATF-deficient (Batf*-/-*) Th17 cells. Western blot analysis showed a marked reduction in XPC protein levels in Batf-/- Th17 cells compared to WT controls **(Figure 2M)**. Quantification of the normalized XPC intensity confirmed a significant decrease in XPC expression in the absence of BATF **(Figure 2N)**. This confirms that BATF is a critical transcriptional regulator of XPC expression during Th17 differentiation, ensuring that sufficient DNA repair capacity is available to support the proper development of these cells.

### The deficiency of XPC impairs the effector program of Th17 cells

Next, we sought to investigate if the impaired IL-17A production by XPC-deficient Th17 cells was related to changes in their effector function. The differentiation and function of Th17 cells rely on the activity of ROR*γ*t, a transcription factor responsible for the expression of lineage signature genes such as *Il17a* and *Il23r* ^1,2,3^. We observed that the reduction of IL-17A production by XPC-deficient Th17 cells was associated with a significantly decreased expression of ROR*γ*t **(Figures 3A-E)**. In addition, these findings go along with a decreased STAT3 phosphorylation, and a decline in the expression of IL-23R but not IL1R1 in the Th17 cells that developed in the absence of XPC **(Figures 3F-N)**.

**Figure 3:**
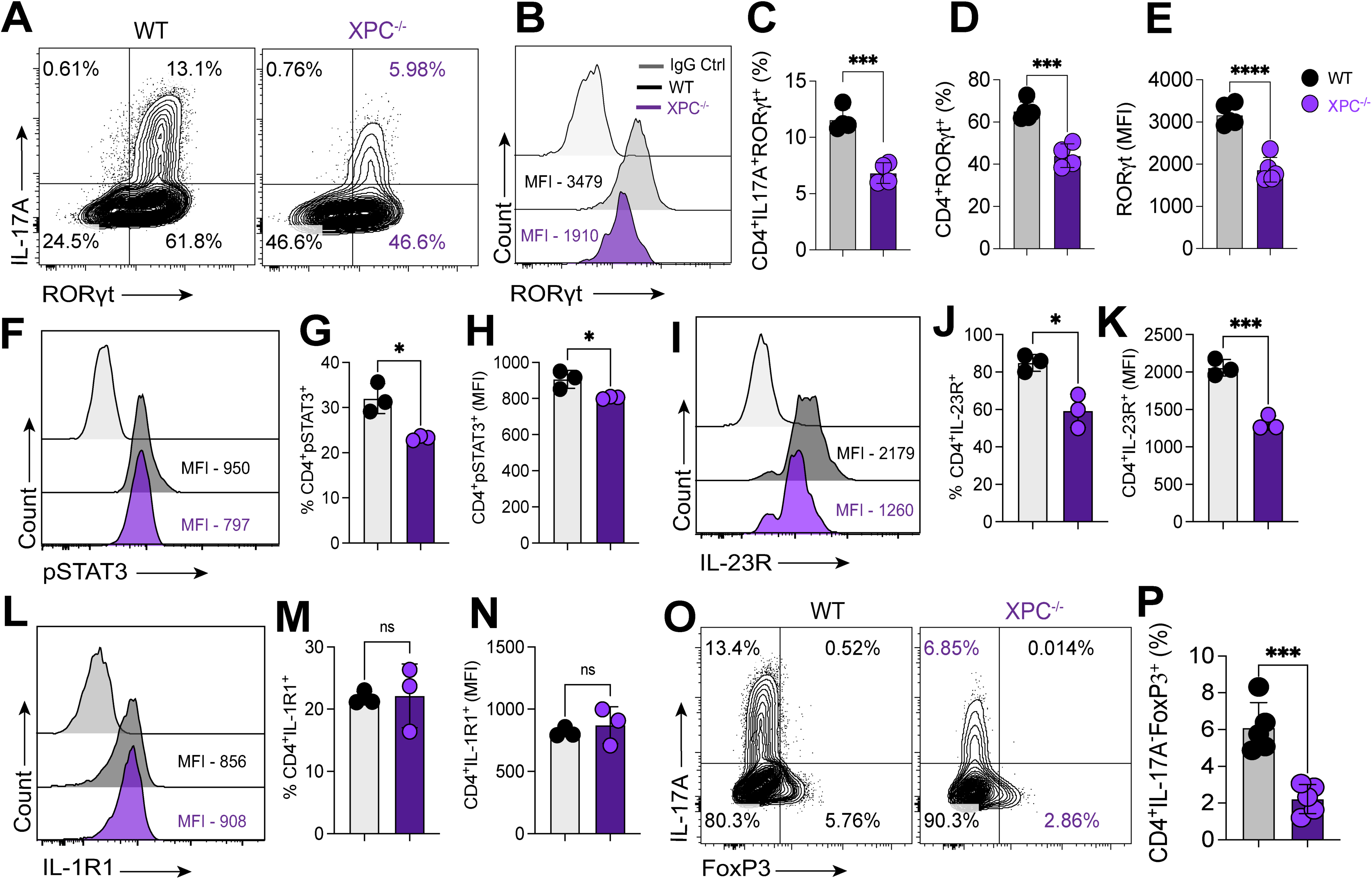
Impaired differentiation and altered expression of key signaling molecules in XPC-deficient Th17 cells. **(A)** Flow cytometry contour plots showing IL-17A and RORγt expression in CD4+ T cells from WT and XPC-/- mice. Percentages of IL-17A+RORγt+ and IL-17A-RORγt+ populations are indicated. **(B)** Flow cytometry histogram comparing RORγt expression in CD4+ T cells from WT and XPC-/- mice. MFI values for each group are shown. **(C)** Quantification of CD4+IL-17A+RORγt+ T cells as a percentage of total CD4+ cells in WT and XPC-/- mice. **(D)** Quantification of CD4+RORγt+ T cells as a percentage of total CD4+ cells in WT and XPC-/- mice. **(E)** RORγt expression (MFI) in CD4+ T cells from WT and XPC-/- mice. **(F)** Flow cytometry histogram showing phosphorylated STAT3 (pSTAT3) expression in CD4+ T cells from WT and XPC-/- mice. MFI values are indicated. **(G)** Percentage of CD4+pSTAT3+ T cells in WT and XPC-/- mice. **(H)** pSTAT3 expression (MFI) in CD4+ T cells from WT and XPC-/- mice. **(I)** Flow cytometry histogram showing IL-23R expression in CD4+ T cells from WT and XPC-/- mice. MFI values are shown for each group. **(J)** Percentage of CD4+IL-23R+ T cells in WT and XPC-/- mice. **(K)** IL-23R expression (MFI) in CD4+ T cells from WT and XPC-/- mice. **(L)** Flow cytometry histogram showing IL-1R1 expression in CD4+ T cells from WT and XPC-/- mice. MFI values are indicated. **(M)** Percentage of CD4+IL-1R1+ T cells in WT and XPC-/- mice. **(N)** IL-1R1 expression (MFI) in CD4+ T cells from WT and XPC-/- mice. **(O)** Flow cytometry contour plots showing IL-17A and FoxP3 expression in CD4+ T cells from WT and XPC-/- mice. Percentages of IL-17A+FoxP3+ and IL-17A-FoxP3+ populations are indicated. **(P)** Quantification of CD4+IL-17A+FoxP3+ T cells as a percentage of total CD4+ cells in WT and XPC-/- mice. Data are representative of at least two independent experiments and are shown as mean ± SEM. *p < 0.05 determined T-test *p < 0.05

The differentiation of Th17 cell and Tregs share a common signaling pathway involving TGF-β. In this context, low concentrations of TGF-β, in conjunction with IL-6 and IL-21, promote the differentiation of Th17 cells, whereas high concentrations inhibit Th17 cells differentiation and instead promote the expression of FOXP3, leading to Treg differentiation. Furthermore, FOXP3 interacts with RORγt and inhibits its ability to bind to the *Il-17a* promoter region (ICHIYAMA et al., 2008a; ZHOU et al., 2008). We investigated if the decreased expression of ROR*γ*t in XPC-deficient Th17 cells could be related to an increase in FoxP3 expression. Surprisingly, we found that even the expression of FoxP3 was downregulated in XPC^-/-^ Th17 cells.

### CD4+ T cells lacking XPC exhibit an inability to differentiate into IL-17A^+^/IFN-γ^+^ producing cells and are incapable of inducing T cell-induced colitis in Rag1^−/−^ mice

To investigate the impact of XPC absence on the function and differentiation of CD4^+^ T cells *in vivo*, we sorted CD4^+^CD45RB^high^ naive T cells from both WT and XPC^−/−^ mice and transferred them into Rag1^−/−^ mice. Subsequently, we monitored the development of colitis. Conversely to Rag1^−/−^ mice receiving WT CD4+ T cells, those that received XPC^−/−^ CD4+ T cells did not exhibit weight loss or colon shortening and displayed reduced colon inflammation. Endoscopy analysis revealed fewer signs of granularity, fibrin, and diarrheic stool in Rag1^−/−^ mice that received XPC-deficient CD4+ T cells **(Figures 4A-F)**.

**Figure 4:**
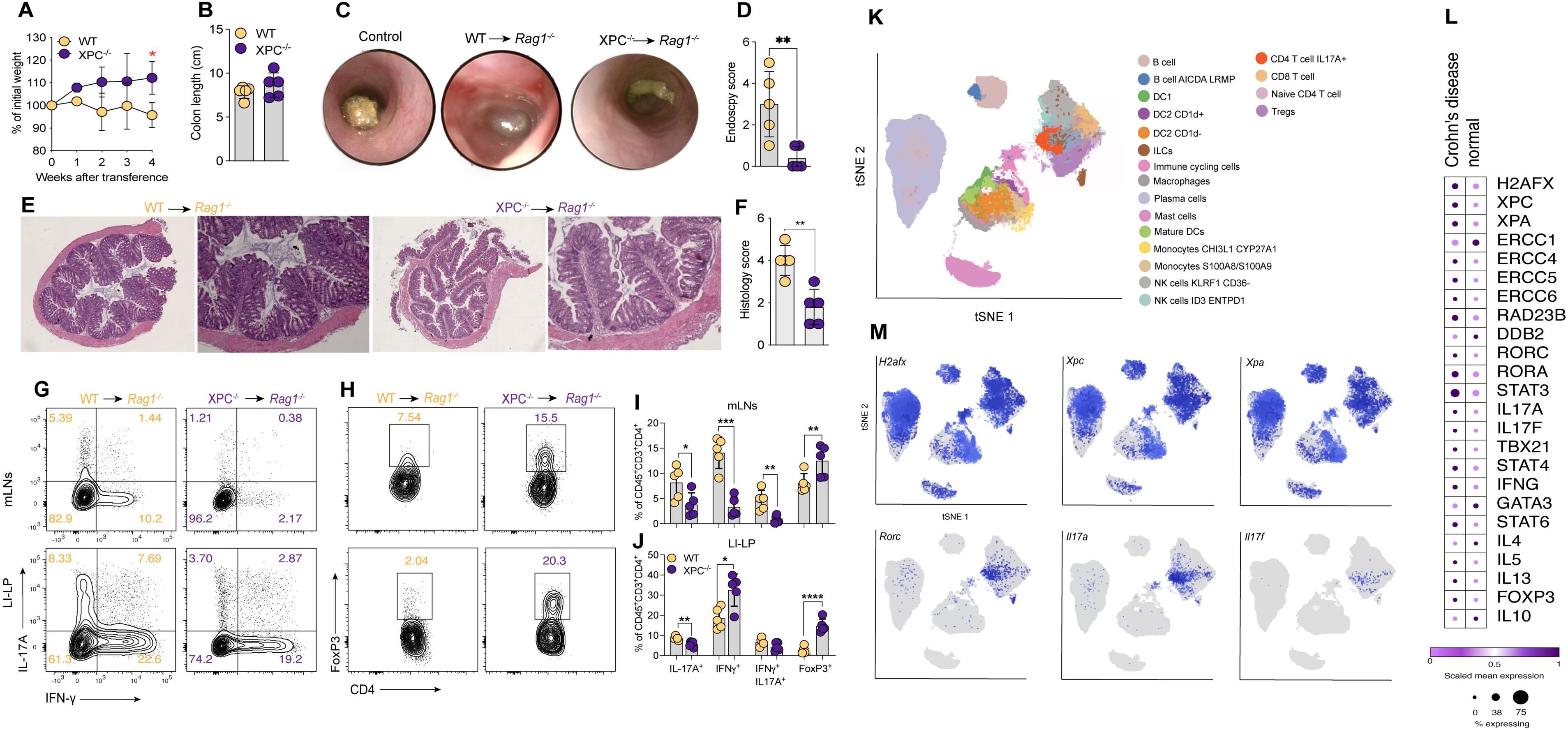
Induction of colitis by CD4+ T cell transfer from WT and XPC-/- mice into Rag1-/- mice, and single-cell RNA-seq analysis of CD45+ cells from Crohn’s disease patients. **(A)** Weight change as a percentage of initial weight in Rag1-/- mice receiving CD4+CD45+RBhigh T cells from WT and XPC-/- mice, monitored over 4 weeks post-transfer. **(B)** Colon length measured at the end of the experiment in Rag1-/- mice receiving CD4+ T cells from WT or XPC-/- mice. **(C)** Representative colonoscopy images showing colon inflammation in control, WT, and XPC-/- cell recipients. **(D)** Endoscopy score quantifying the severity of intestinal inflammation in Rag1-/- mice receiving WT or XPC-/- CD4+ T cells. **(E)** Representative histological sections of colonic tissue from Rag1-/- mice after T cell transfer from WT and XPC-/- donors. **(F)** Histology score quantifying intestinal inflammation in Rag1-/- mice based on colonic tissue sections. **(G)** Flow cytometry plots showing IFN-γ and IL-17A production by CD4+ T cells from mesenteric lymph nodes (mLNs) and lamina propria (LI-LP) in Rag1-/- mice receiving WT or XPC-/- CD4+ T cells. **(H)** Flow cytometry plots showing FoxP3 expression in CD4+ T cells from mLNs and LI-LP of Rag1-/- mice receiving WT or XPC-/- CD4+ T cells. **(I)** Quantification of CD4+CD45+ T cell subsets (IL-17A+, IFN-γ+, and FoxP3+) from mLNs in Rag1-/- mice receiving WT or XPC-/- CD4+ T cells. **(J)** Quantification of CD4+CD45+ T cell subsets (IL-17A+, IFN-γ+, FoxP3+) from LI-LP in Rag1-/- mice receiving WT or XPC-/- CD4+ T cells.(K) t-SNE analysis of immune cell populations from inflamed colonic tissue of Crohn’s disease patients, identifying various immune cell clusters, including Th17, Th1, and Treg cells. **(L)** Heatmap showing scaled expression of DNA repair genes (Xpc, Ercc1, etc.), transcription factors (RORγt, STAT3), and cytokines (IL-17A, IFN-γ) in colonic tissues from Crohn’s disease patients, comparing inflamed and normal conditions. **(M)** t-SNE plots showing gene expression (H2afx, Xpc, Xpa, Rorc, Il17a) in various immune cell populations from colonic tissue of Crohn’s disease patients. Data available at Single Cell portal at the Broad Institute (SCP1884) and obtained from the study of Kong et al., (2023). Data are representative of at least two independent experiments and are shown as mean ± SEM. *p < 0.05 determined by one-way ANOVA followed by Tukey’s post hoc test or T-test *p < 0.05.

Subsequently, we assessed cytokine production by WT and XPC-deficient CD4+ T cells in the mesenteric lymph nodes (mLNs) and colonic lamina propria (c-LP) of Rag1^−/−^ mice four weeks post-T cell transfer. The frequency of TCR-β^+^CD4^+^IL17A^+^ and TCR-β^+^CD4^+^IL17A^+^IFN-γ^+^ producing cells were significantly diminished in mice transferred with XPC-deficient CD4+ T cells compared to those receiving WT CD4+ T cells **(Figures 4G-J)**. Furthermore, the frequency of TCR-β^+^CD4^+^FOXP3^+^ T cells was increased in mice transferred with XPC-deficient CD4+ T cells compared to Rag1^−/−^ mice receiving WT ones **(Figures 4H, 4I, 4J)**.

Next, we analyzed single-cell data obtained from Crohn’s disease (CD) patients, accessible through the Single Cell portal at the Broad Institute (SCP1884) and obtained from the study of Kong et al. (2023) ^23^, in order to understand if Th17 cells from CD patients express genes related to DNA damage and the NER DNA repair pathway including XPC. In this dataset, CD45^+^ cells were isolated from colon biopsies of both healthy controls and individuals with CD, and then subjected to single-cell sequencing. CD45^+^ cells derived from CD patients displayed heightened expression levels of genes associated with DNA repair, including *XPA*, *XPC*, *ERCC4*, *ERCC6* and *RAD23B*. Th17 cells from in CD patients exhibited increased expression of XPC alongside with the classical genes of its signature, *Rorc*, *Stat3*, *Il17a* and *Il17f*, suggesting a potential role for XPC in the differentiation of human Th17 cells during the course of CD **(Figures 4K-M)**.

### The deficiency of XPC alters the expression of genes associated with DNA repair pathways in Th17

To unravel the mechanism by which XPC influences Th17 cell differentiation, we conducted bulk RNA-seq on both WT and XPC-deficient Th17 cells. Remarkably, our analysis identified 1758 genes that were significantly downregulated and 1447 genes that were upregulated in XPC-deficient Th17 cells **(Figure 5A)**. Gene ontology analysis unveiled an enrichment of genes associated with various pathways, prominently including DNA repair and DNA replication which is important for DNA processing and repair **(Figure 5B)**. Based on the log2FC, we rank the top different expressed genes (DEGs), and we observed that genes such as *Exo1* involved in the BER DNA repair pathway was significant downregulated and the gene *Txnip* that encodes the protein Thioredoxin-interacting protein (TXNIP) and is involved in the inhibition of antioxidant response ^24^ was significant upregulated in the absence of XPC in Th17 cells (**Figure 5C)**. The gene enrichment network illustrated interconnected and enriched genes from pathways such as BER, NER, and mismatch repair, intertwined with pathways related to cell cycle, cellular senescence, and metabolic processes **(Figure 5D)**.

**Figure 5:**
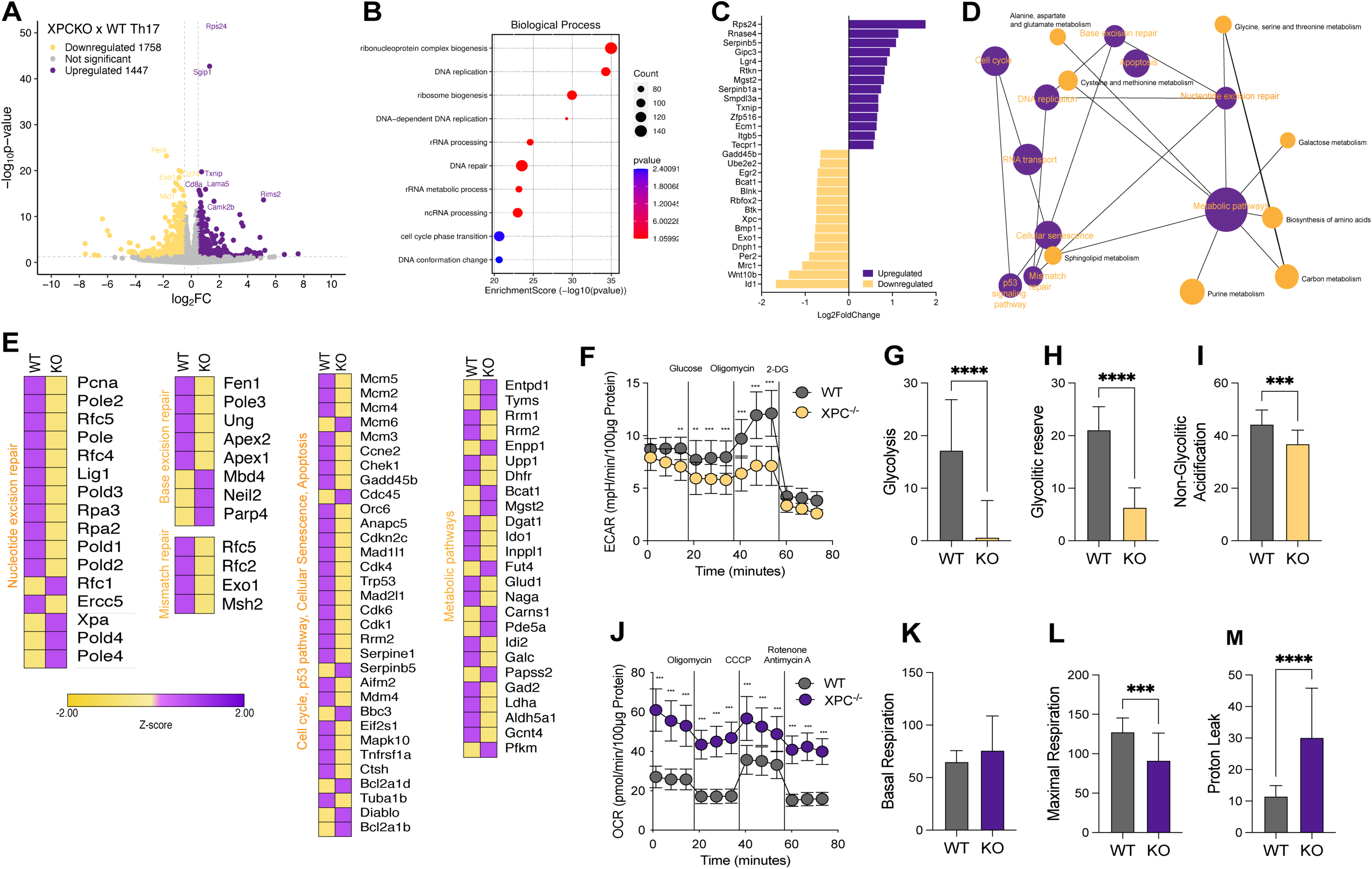
Transcriptomic and metabolic differences between WT and XPC-/- Th17 cells. **(A)** Volcano plot showing differentially expressed genes (DEGs) between XPC-/- and WT Th17 cells. Genes significantly upregulated in XPC-/- cells are highlighted in yellow, and those downregulated are in purple. **(B)** Gene Ontology (GO) enrichment analysis of DEGs, showing biological processes significantly enriched in XPC-/- Th17 cells. **(C)** Bar plot of top differentially expressed genes based on log2 fold-change (log2FC) between XPC-/- and WT Th17 cells. **(D)** Network map showing the interaction between metabolic and DNA repair pathways enriched in XPC-/- Th17 cells, with highlighted pathways involved in cell cycle, DNA repair, and metabolism. **(E)** Heatmaps displaying the Z-scores of genes involved in nucleotide excision repair, base excision repair, mismatch repair, cell cycle, and metabolic pathways in WT and XPC-/- Th17 cells. **(F)** Extracellular acidification rate (ECAR) measurements over time, comparing glycolytic activity between WT and XPC-/- Th17 cells after sequential treatment with glucose, oligomycin, and 2-deoxy-D-glucose (2-DG). **(G)** Quantification of glycolysis rates in WT and XPC-/- Th17 cells. **(H)** Quantification of glycolytic reserve in WT and XPC-/- Th17 cells. **(I)** Quantification of non-glycolytic acidification in WT and XPC-/- Th17 cells. **(J)** Oxygen consumption rate (OCR) measurements over time, comparing mitochondrial respiration between WT and XPC-/- Th17 cells after sequential treatment with oligomycin, CCCP, and rotenone/antimycin A. **(K)** Basal respiration rates in WT and XPC-/- Th17 cells. **(L)** Maximal respiration rates in WT and XPC-/- Th17 cells. **(M)** Proton leak measurements in WT and XPC-/- Th17 cells, comparing mitochondrial efficiency. Data are representative of at least two independent experiments and are shown as mean ± SEM. *p < 0.05 determined T-test *p < 0.05

Upon detailed examination of the altered genes within these pathways, a notable reduction in the expression of several genes associated with the NER pathway was observed, including *Pcna*, *Lig1*, *Pole*, and *Ercc5* **(Figure 5E)**. Furthermore, downregulation extended to genes linked with the cell cycle, p53 pathway, and cellular senescence. Conversely, genes associated with resistance to apoptosis, such as *Bcl2a1d* and *Bcl2a1b*, exhibited upregulation in the absence of XPC **(Supplementary Figure 2A)**.

The absence of XPC in Th17 cells is associated with the downregulation of genes associated with various metabolic pathways, such as Dgat1 and Ldha, while showing an upregulation of genes like *Mgdt2* and *Pd5a*, which are associated with oxidative stress **(Figure 5E)** and **(Supplementary Figure 2B-G)**. In line with these, FACS analysis indicated decreased levels of the apoptosis marker Annexin V and a reduction in the proliferation marker Ki67 in XPC-deficient Th17 cells **(Supplementary Figure 2H-M)**. Also, we found that XPC-deficient Th17 cells display a reduction in Extracellular Acidification Rate (ECAR), an indicative of diminished glycolysis **(Figures 5F-I)**, and an elevation in Oxygen Consumption Rate (OCR), suggesting heightened mitochondrial respiration **(Figures 5J, 5K, 5L, 5M)**.

### XPC-deficient Th17 cells accumulates oxidative DNA damage and interact with OGG1

Given the crucial role of XPC in DNA repair, we hypothesized that its absence might lead to increased DNA damage in Th17 cells, resulting in genomic instability and compromising their differentiation.

In order to understand if Th17 cells accumulates DNA damage in the absence of XPC, we cultured WT and naïve XPC-deficient CD4+ T cells under Th17 cells polarizing conditions and measured DNA damage by the expression of the phosphorylated γH2AX and comet assay. Our findings indicated elevated levels of γH2AX and increased DNA damage in XPC-deficient Th17 cells when compared to WT cells **(Figures 6A-C)**. In addition, alkaline comet assay indicates increased DNA breaks lesions as well as increased sensitivity of formamidopyrimidine DNA glycosylase (FPG) sites, which suggest increased oxidized bases in XPC-deficient Th17 cells **(Figures 6D-E)**. Moreover, BATF-deficient Th17 cells exhibited a phenotype resembling that of XPC-deficient Th17 cells, characterized by elevated levels of mitochondrial ROS and increased DNA damage **(Supplementary Figure 3A-D).**

**Figure 6:**
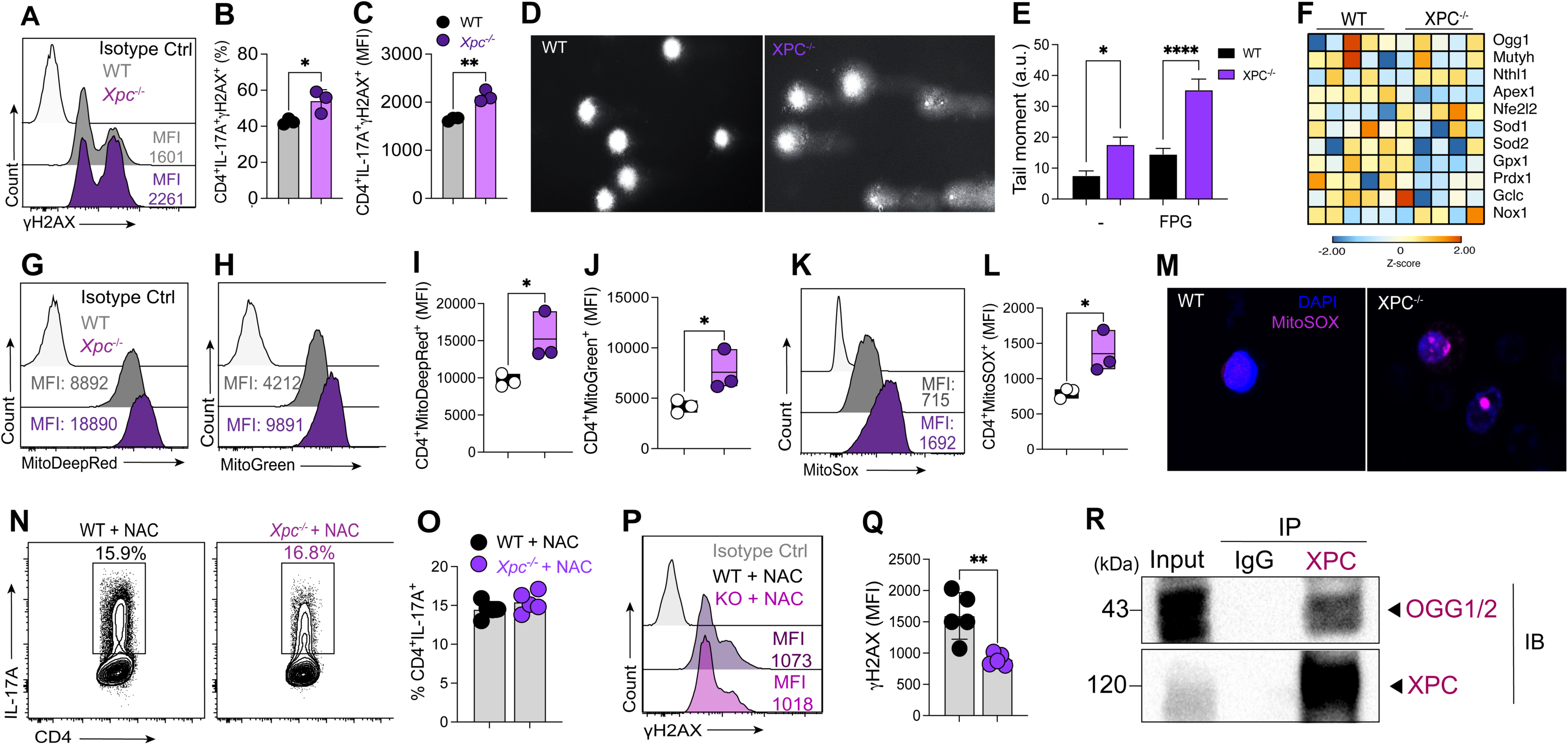
Mitochondrial oxidative stress and DNA damage in XPC-/- Th17 cells. **(A)** Flow cytometry histogram showing γH2AX expression in CD4+IL-17A+ cells from WT and XPC-/- conditions. MFI values are indicated. **(B)** Quantification of CD4+IL- 17A+γH2AX+ cells as a percentage of total CD4+ cells. **(C)** γH2AX expression (MFI) in CD4+IL-17A+ cells from WT and XPC-/- conditions. **(D)** Representative comet assay images showing DNA damage in CD4+ cells from WT and XPC-/- conditions. **(E)** Quantification of tail moment in the comet assay for CD4+ cells treated with or without FPG to detect oxidative DNA damage. **(F)** Heatmap of genes involved in oxidative stress and antioxidant responses, comparing WT and XPC-/- Th17 cells. **(G-H)** Flow cytometry histograms showing MitoDeepRed and MitoGreen staining in CD4+ cells, indicating mitochondrial mass and membrane potential. **(I-J)** Quantification of MitoDeepRed+ and MitoGreen+ CD4+ cells (MFI). **(K)** Flow cytometry histogram showing MitoSOX staining in CD4+ cells, indicating mitochondrial superoxide levels. **(L)** Quantification of MitoSOX+ CD4+ cells (MFI). **(M)** Representative microscopy images showing MitoSOX staining in CD4+ cells, indicating mitochondrial oxidative stress. **(N-O)** Flow cytometry plots and quantification of IL-17A expression in CD4+ cells treated with N-acetylcysteine (NAC), a ROS scavenger. **(P-Q)** Flow cytometry histograms and quantification of γH2AX expression (MFI) in CD4+IL-17A+ cells treated with NAC. **(R)** Co-immunoprecipitation and immunoblot analysis of the interaction between XPC and OGG1/2 in CD4+ cells. Data are representative of at least two independent experiments and are shown as mean ± SEM. *p < 0.05 determined by one-way ANOVA followed by Tukey’s post hoc test or T-test *p < 0.05.

**Figure 7:**
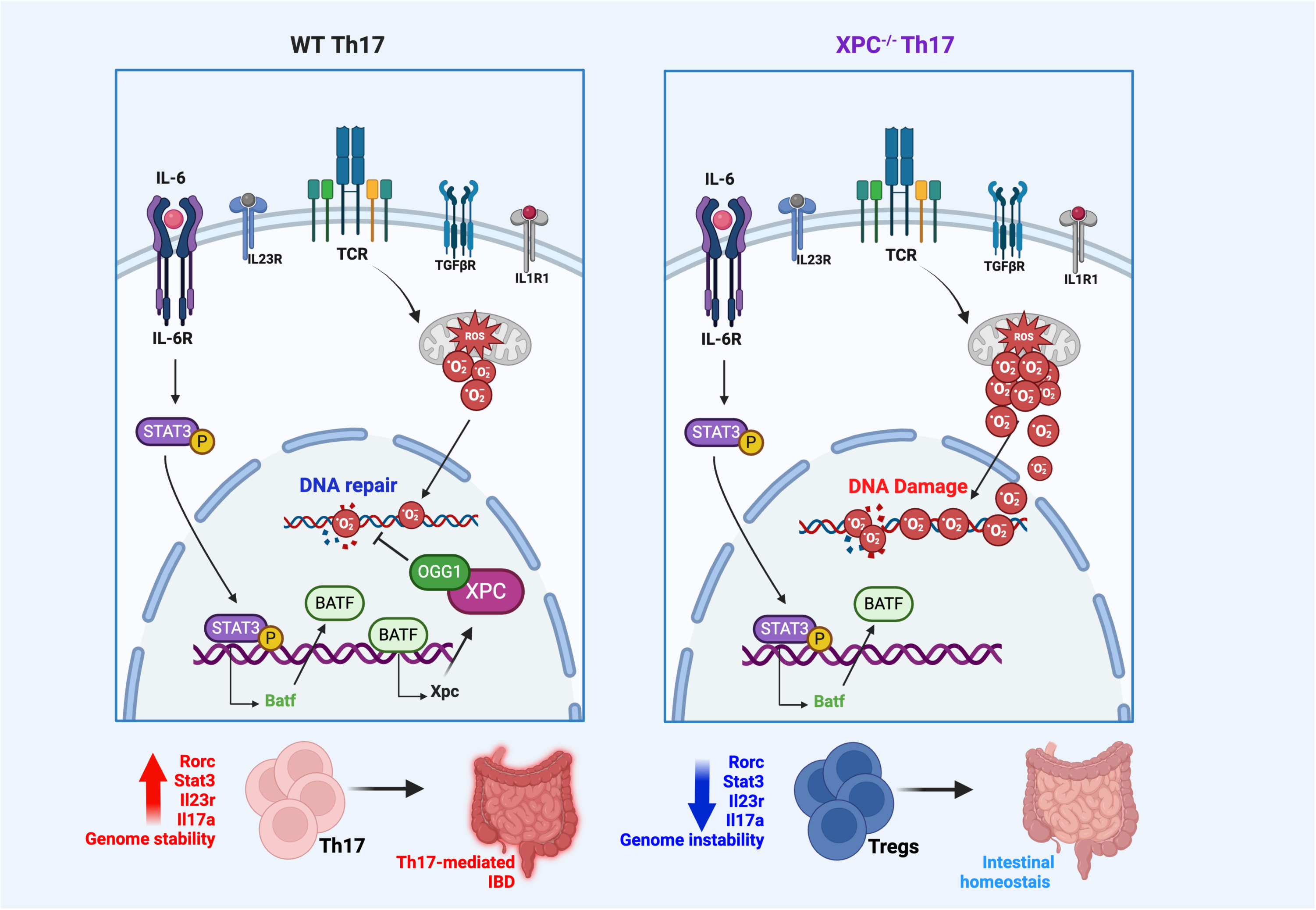
Graphical abstract illustrating the role of XPC in maintaining genome stability during Th17 differentiation and its impact on intestinal inflammation.. Left panel (WT Th17): In WT Th17 cells, signaling via IL-6R and IL-23R leads to STAT3 phosphorylation, promoting BATF and Rorc transcription. ROS generated during activation are mitigated by the DNA repair machinery, including XPC and OGG1, preserving genome stability. This leads to the robust differentiation of Th17 cells, characterized by high expression of Il17a, Il23r, and Stat3, which are involved in promoting inflammation and driving Th17-mediated intestinal inflammation (IBD). Right panel (XPC-/- Th17): In XPC-/- Th17 cells, the absence of functional XPC results in impaired DNA repair in response to ROS-induced damage. This leads to genome instability, which interferes with Th17 differentiation. Instead of promoting inflammation, XPC-deficient cells exhibit reduced expression of Il17a, Il23r, and Stat3, with a skewing towards Treg differentiation. This shift towards Tregs contributes to maintaining intestinal homeostasis and reduces inflammation. Bottom panel: The differences between WT and XPC-/- Th17 cells culminate in distinct outcomes: WT Th17 cells drive intestinal inflammation and IBD, while XPC-/- Th17 cells promote Treg differentiation, supporting intestinal homeostasis.

Our RNA-seq data reveals a decrease in the expression of genes such as *Ogg1*, *Sod1*, *Sod2*, and *Gpx1*, which are involved in the repair of oxidized bases and antioxidant response. XPC-deficient Th17 cells also upregulate *Nox1*, the gene encoding NADPH enzyme participating in the generation of ROS and NFe2L2, the gene that encodes the transcription factor NRF2 that is important for the regulation of genes with antioxidant functions **(Figure 6F)**. We found that XPC-deficient cells exhibit an increased mitochondrial membrane potential and elevated mitochondrial ROS levels, suggesting the possibility of mitochondrial dysfunction and oxidative stress due to defects in DNA repair **(Figures 6G-M)**.

Due to the observation of increased oxidative stress and DNA damage in XPC-deficient Th17, we hypothesize that the inhibition ROS could rescue the accumulation of DNA damage and therefore IL-17A production. In order to test this hypothesis, we cultured naïve WT and XPC-deficient CD4+ T cells under Th17 cells polarizing conditions in the presence of N-Acetylcysteine (NAC). NAC treatment was able to rescue IL-17A production in XPC^-/-^ Th17 cells **(Figure 6N-O)**. Moreover, NAC treatment decreased γH2AX expression in XPC-deficient Th17 cells which suggests that NAC was able to decrease oxidative stress and DNA damage and therefore rescue Th17 cells differentiation **(Figure 6P-Q)**.

Since we found that XPC-deficient Th17 cells are more sensitive to increased ROS levels which leads to DNA damage and genomic instability, we hypothesized that XPC could interact with DNA repair proteins that are involved in the repair of oxidized DNA damage. In this sense, OGG1 is the key enzyme responsible for removing 8-oxoguanine (8-oxoG), a mutagenic byproduct of exposure to ROS. Therefore, we investigated whether XPC and OGG1 can interact during the development of Th17 cells. We performed *in vitro* Th17 cells differentiation and harvested the cells at 96h of culture and performed co-immunoprecipitation analysis. Indeed, we found that XPC co-immunoprecipitated with OGG1 **(Figure 6R)**.

Taken together, our results suggest that the absence of XPC leads to increased DNA damage in Th17 cells, impairing their differentiation due to genomic instability. Treatment with NAC rescues IL-17A production and decreases DNA damage in XPC-deficient CD4+ T cells, suggesting a potential therapeutic avenue. Additionally, the interaction between XPC and OGG1, highlights XPC’s role in repairing oxidized DNA damage during Th17 cell differentiation.

## Discussion

T cells are frequently exposed to harmful factors that can threaten genomic stability. These can lead to alterations in their differentiation that can impact autoimmunity^25^. ROS are commonly regarded as detrimental byproducts of cellular metabolism, causing damage to nucleic acids ^10^. In autoimmune conditions, T cells exhibit abnormal expression of DNA damage response (DDR) genes and proteins, coupled with oxidative stress. These factors have been linked to changes in the differentiation of cells into immunogenic subsets and/or an elevated rate of apoptosis ^26,27^. The increased DNA damage observed in T cells during autoimmunity could also be associated with a compromised DNA repair^21^. However, it remains unknown how T cells use the DNA repair machinery to remove DNA damage. Here, we define the DNA repair protein XPC as a positive regulator of Th17 cells differentiation and function by protecting them from oxidative DNA damage. The deficiency of XPC leads to compromised Th17 cell differentiation and function, impacting inflammatory responses *in vivo*. Additionally, we found that the transcription factor BATF promotes XPC expression in Th17 cells, which suggests a potential functional relationship between BATF and XPC in maintaining DNA repair machinery and genomic integrity in Th17 cells.

XPC plays an essential role in initiating GG-NER^28^. Individuals with inherited mutations in the XPC and other genes of the XP family, including XP-A, XP-B, XP-D, XP-F, XP-G, XP-V, develop a rare disorder named Xeroderma pigmentosum (XP) characterized by heightened vulnerability to skin cancer, attributed to impaired removal of genotoxic dipyrimidine photoproducts induced by solar UV radiation through GG-NER ^29^. Growing evidence suggests that XP patients have defects in their immune system, which could explain the vulnerability to skin cancer and other conditions observed in these patients. XP patients have reduced numbers of CD4+ T cells as well as reduced NK lytic activity ^30^. In addition, T cells from XP patients exhibited markedly reduced levels of the antiapoptotic Bcl-2 and CD95 molecule compared to non-XP T cells, resulting in the highest Bax/Bcl-2 ratio for XP T-cells suggesting a susceptibility to undergo apoptosis ^31^. On the other hand, some XP patients are susceptible to urinary infections and also display bilateral pneumonia with bacterial and fungal infection ^32^. In our study, we found that Th17 cells that lack XPC display a compromised expression of the main identity proteins RORγt, STAT3 and IL-23R, therefore leading to an impaired function which suggest that XP patients could be more susceptible to these infecting agents due to an impaired Th17 cell differentiation. Th17 cells play a protective role against skin cancers such as squamous cell carcinoma and melanoma, while Treg cells have a pro-tumor function in various types of cancer. Our findings, which show that XPC-deficient mice have an impaired Th17 response along with an increase in Treg cells, may help explain the higher susceptibility to cancer in XP patients. It is possible that the elevated Treg cells within tumors contribute to tumor growth, suggesting that this imbalance between Th17 and Treg cells could be a factor in the increased risk of carcinogenesis in these patients ^33–35^.

The expression of XPC is regulated by a variety of transcription factors that ensure proper DNA repair and genomic stability in different cell types. In Th17 cells, we identified BATF, RORγt, and IRF4 as key regulators of Xpc expression. Among these, BATF emerged as a critical player, with BATF-deficient Th17 cells exhibiting a similar phenotype to XPC deficiency, characterized by increased mitochondrial ROS, DNA damage, and reduced XPC expression. These findings suggest a potential BATF-XPC pathway that plays a central role in DNA repair and maintaining genomic integrity in Th17 cells. BATF is a well-established transcription factor essential for Th17 cell identity and differentiation. BATF-deficient mice show impaired Th17 differentiation and dysregulation of key cytokine genes. BATF directly binds to conserved regulatory elements within the IL-17A/F locus, as well as to the promoters of Il17a, Il21, and Il22, further underscoring its importance in Th17 cell function.^48^. Nevertheless, the deficiency of BATF in Th17 cells is associated with a less enrichment of genes related to DNA repair pathways ^49^. Therefore, the identified direct interaction between XPC and BATF further suggests a potential enhancement of BATF-mediated transcriptional functions facilitated by XPC in Th17 cells.

After identifying a helix-distorting lesion, either XPC or RNA polymerase II initiates NER, involving TFIIH to unwind DNA, XPA stabilization, and collaboration with XPF/ERCC1 for incision. Repair includes coordinated synthesis, further incision by XPG, and nick repair by DNA ligases ^37–39^. The findings from XPC-deficient Th17 cells reveal a significant impact on DNA repair pathways, particularly the NER pathway. Additionally, downregulation extends to genes related to BER and the mismatch repair pathway, along with those associated with the cell cycle, p53 pathway, and cellular senescence. This comprehensive downregulation could explain the compromised differentiation and function of XPC-deficient Th17 cells due to impaired DNA repair. These results suggest that XPC may have an essential role in preserving genomic stability in Th17 cells by ensuring effective DNA repair.

In Th17 cells, the deficiency of STIM-1 and MINK1 is associated with the accumulation of ROS, DNA damage, cell death and genomic instability ^13,14^. A similar phenotype was observed in XPC-deficient Th17 cells with increased DNA damage, oxidative stress and altered cellular metabolism. ROS, including superoxide radicals and hydrogen peroxide, can react with DNA bases. These modifications can disrupt the typical structure of DNA, potentially causing mutations and compromising genomic integrity ^40,41^ When DNA bases undergo oxidation, it can interfere with transcription processes. Oxidized bases can halt the progression of RNA polymerase during transcription, leading to transcriptional errors or even transcriptional arrest ^42^. Nevertheless, fibroblasts exhibiting reduced XPC protein expression undergo a metabolic transition from oxidative phosphorylation (OXPHOS) to glycolysis, associated with increased nuclear DNA damage and the generation of oxidants through NOX-1 ^17^. These results align with our findings that XPC-deficient Th17 cells show increased NOX-1 expression and metabolic changes, including reduced glycolysis and increased OXPHOS. The altered mitochondrial membrane potential, higher mitochondrial ROS levels, and shifts in metabolism, characterized by decreased glycolysis and enhanced mitochondrial respiration, suggest an extensive metabolic shift and mitochondrial dysfunction. These changes, along with increased DNA damage and oxidized bases, may contribute to genome instability and impaired differentiation and function of Th17 cells lacking XPC.

The repair of oxidized bases involves intricate pathways such as BER^16^ and NER^19^. XPC-deficient Th17 cells downregulate genes involved in the repair of oxidized bases such as Ogg1, Mutyh, Apex1, Exo1, as well as genes involved in antioxidant effects such as Sod1, Sod2 and Gpx1 which suggest that XPC is not only important for the regulation of NER DNA repair pathways but also for other pathways such as the BER and mismatch pathways as well as the generation of antioxidant responses in Th17 cells. These results are consistent with previous findings that XPC-deficient fibroblasts exhibited decreased levels of endogenous APE1 and OGG1 mRNA.^43^. In addition, keratinocytes and fibroblasts that lack XPC display a reduction in the repair efficiency of 8-OH-Gua. Thus, the XPC-HR23B complex serves as a cofactor in the base excision repair process of 8-OH-Gua, enhancing the activity of the DNA glycosylase, OGG1 ^18^. In line with this, we have identified a physical interaction between XPC and OGG1 in Th17 cells, suggesting a potential link between these proteins. This interaction might enhance XPC’s role in the complex processes of base excision repair. The downregulation of genes involved in repairing oxidized bases and antioxidant defenses in XPC-deficient Th17 cells implies that XPC is crucial for coordinating repair mechanisms, including those involving OGG1. This underscores the importance of XPC in maintaining genomic stability and cellular function under oxidative stress within Th17 cells..

Besides its function as a DNA damage sensor, XPC also functions as a transcription regulator by interacting with E2F1 and promotes its binding to its DNA element ^44^. The transcription factor E2F1 is a central player involved in cell cycle progression, DNA-damage response, and apoptosis ^45^ and its target genes include Ogg1 and Mutyh. In addition, E2F1 is a protective factor against oxidative stress ^46^. In addition, XPC is required for efficient RNA synthesis, optimal DNA demethylation, and histone posttranslational modifications (H3K4/H3K9 methylation, H3K9/14 acetylation) ^47^. Thus, the observed decrease in DNA repair and antioxidant response gene expression in Th17 cells lacking XPC suggests that XPC’s transcriptional regulatory role plays a crucial part in modulating the cellular response to oxidative damage.

In summary, our study reveals the role of XPC in maintaining genomic stability and influencing Th17 cell differentiation. Th17 cells upregulating XPC exhibit lower oxidative DNA damage levels. XPC deficiency impairs Th17 differentiation, evidenced by reduced IL-17A production and altered expression of key Th17 markers. *In vivo* experiments show that XPC-deficient Th17 cells fail to induce T cell-induced colitis. Transcriptomic analysis highlights the impact of XPC deficiency on DNA repair, metabolism, and oxidative stress pathways, indicating intricate connections between these processes.

This comprehensive exploration of XPC’s role in Th17 cells not only advances our understanding of DNA repair mechanisms in T cells but also highlights the intricate connections between DNA repair, cellular metabolism, and immune responses. The impairment in the immune system may partially contribute to the high frequency of skin tumors in XP patients. The implications of our study extend to broader contexts, including autoimmune diseases and bacterial and fungal infections, where Th17 cells play a pivotal role. Targeting the XPC pathway could offer new avenues for therapeutic interventions in conditions associated with Th17 cell dysregulation.

## Methods details

### Animals

Male wild-type C57BL/6, XPC^-/-^ and BATF^-/-^ mice aged 6-8 weeks and weighing 20-25 grams were used. All procedures were performed in accordance with the principles proposed by the Brazilian College of Animal Experimentation (COBEA) and approved by the Ethics Committee on Animal Experimentation of the University of São Paulo (CEUA ICB: 35881860349).

### In vitro T helper cells polarization

In vitro T helper cells polarization was performed by isolating naïve CD4+ T cells from the spleen and lymph nodes of WT and XPC-/- mice using microbeads coated with anti-CD4 (clone L3T4, cat 130-117-043, Miltenyi Biotec, Bergisch Gladbach, Germany). The cells were then stained with fluorochrome-conjugated anti-CD4, anti-CD44, and anti-CD62L antibodies, and CD4+CD44lowCD62Lhigh cells were isolated by sorting (FACS Aria III sorter, BD Biosciences). For the differentiation, the CD4 T cells obtained from the sorting were placed in 48-well plates coated with anti-CD3 (2 ug/mL – BD Bioscience Cat. 553058, Clone – 145-2C11) anti-CD28 (1 ug/mL - BD Bioscience Cat. 553294, Clone: 37.51 RUO), and subsequently stimulated with a cocktail of cytokines and neutralizing antibodies. Th17: 2.5 ng/mL of TGF-beta (R&D – Cat. 7666MB), 50 ng/mL of IL-6 (Peprotech – Cat. 216-16), 50 ng/mL of IL-23 (R&D, Cat. 1887-ML), 20 ng/mL of IL-1beta (Peprotech, Cat. 211-11B) and 1 ug/mL of anti-IFNg (BD Bioscience, Cat. 554408), anti-IL-4 (BD Bioscience, Cat. 554432, Clone: 11B11) and anti-IL-2 (BD Bioscience, Cat. 554424, Clone: JES6-1A12). Th1: 20 ng/mL of IL-12 (Cat. 212-12, Peprotech) and 1 ug/mL anti-IL-4 (Cat. 554432, BD Bioscience). Th2: 10 ng/mL of IL-4 (Cat. 214-14, Peprotech) and 1 ug/mL of anti-IFNg (BD Bioscience, Cat. 554408). iTreg: 10 ng/mL TGF-beta (R&D – Cat. 7666MB) and 1 ug/mL of anti-IFNg (BD Bioscience, Cat. 554408).

### Flow cytometry (FACS)

Quantitative and phenotypic analysis of cells differentiated in vitro or isolated from lymph nodes, spleen, or intestinal lamina propria was performed by flow cytometry. Briefly, after being isolated, cells were counted and evaluated for viability with trypan blue. Subsequently, the cells were plated in a 96-well plate and stimulated with phorbol myristate acetate (PMA) (50 ng/ml; Sigma Aldrich) and ionomycin (500 ng/ml; Sigma Aldrich), and brefeldin A (transport inhibitor of golgi - 1.5 μl/ml; BD Biosciences), for 4 hours in an oven at 37°C (5% CO2 and 8% humidity). The number of cells used were 5×10^5/cells per well for experiments derived from in vitro culture of lymphocytes or 2×10^6/cells per well for in vivo experiments. Then, the cells were washed twice with PBS 1x and centrifuged at 450 g for 5 minutes and incubated for 15 minutes with Fc block (1:200) to block Fc receptors, and subsequently washed with PBS 1x, centrifuged at 450 g per 5 and stained with cell viability marker (Fixable viability dye; 1:1000; eBioscience) and anti-CD4 (1:200; BD Biosciences) for 10 minutes at room temperature (Table II). Then, the cells were centrifuged and fixed using the kit’s fixative (transcription factor staining buffer set, eBioscience) for 60 minutes at room temperature, with subsequent centrifugation. The next step was intracellular labeling with antibodies listed in Table I for 60 minutes. Briefly, the antibodies were diluted in the kit’s permeabilizer, and the cells were incubated with permeabilization buffer containing the antibody mix for 60 minutes at 4°C. After this step, the cells were centrifuged again and resuspended in PBS for acquisition by flow cytometry (FACS Canto II, BD Biosciences). In this step, between 100,000 and 250,000 events were acquired according to the experiment. Sample analyses were performed using the FlowJo X software (BD Biosciences).

### Real-time PCR

To assess gene expression of Th17 T cell cultures, cells were harvested, and RNA was extracted using the RNA Extraction Kit (Promega) according to the manufacturer’s instructions. After extraction, samples were quantified using a nanodrop (ThermoFisher) and 500 ng of RNA were converted to cDNA using the High-Capacity cDNA Reverse Transcription Kit (ThermFisher), according to the manufacturer’s instructions. Real-time PCR reactions were performed using cDNA, primers, and the Sybr Green Master Mix containing reaction products such as nucleotides and polymerase. The reactions were performed with the StepOnePlus device (ThermoFisher). The results were analyzed using the cycle threshold (CT), with the data normalized by the Gapdh expression and the difference between the groups calculated using the 2-ΔΔCT method. The sequences of the primers used are described in the table below. None of the primers used showed amplification in the group without sample (blank) or more than one peak in the melting curve. In addition, all primers showed an efficiency greater than 90%.

### RNA-seq analysis

Total RNA extraction from five independent biological replicates of each Th17 WT and KO cells was performed using TRIzol reagent (Life Technologies, Carlsbad, CA, United States) following manufacturer instructions. After, RNA samples were treated with DNase I (1 U/µg of RNA) (Thermo Scientific, Lithuania, EU) at 37°C for 15 min. The absence of DNA contamination was determined from the A260/A280 ratio using a spectrophotometer Nanodrop ND1000 (Thermo Scientific, United States). RNA integrity was evaluated using an Agilent 2,100 Bioanalyzer and a Pico Agilent RNA 6000 kit (Agilent Technologies, Santa Clara, CA, United States). rRNA depletion was performed using Ribo-Zero Human/rat/mice Ribo-Zero plus rRNA Depletion Kit (Illumina). Library preparation was performed using 1 µg of rRNA-depleted total RNA using Truseq Stranded Total RNA LT Sample Prep Gold kit, without molecular barcodes. Sample preparation followed preparation using the NovaSeq 6000 S4 Reagent Kit (Illumina) according to manufacturer’s instructions for RNA sequencing submission. The sequencing was performed with paired-ends (100 bp) using the Illumina Novaseq 6,000 Platform in the Center for Cell-Based Therapy Facility at Ribeirao Preto Medical School Ribeirão Preto. The raw data was deposited at Sequence read archive - SRA under the accession code: PRJNA1160718.

### Western blot

To evaluate XPC protein expression, samples were collected in RIPA buffer (Sigma Aldrich), homogenized with protease, centrifuged, and the supernatant collected. Then, protein quantification was performed using the Bicinchoninic acid protein assay (BCA; Sigma Aldrich). For the experiments with samples of CD4 T cells, 30 μg of proteins were used. Sample preparation was performed with 2x Laemmli sample buffer (Bio-rad), and heated for 10 minutes at 95°C for protein denaturation. Subsequently, the samples were applied to a 12% gradient polyacrylamide gel (Bio-rad) and, after separation, were transferred to 0.2 μm nitrocellulose membranes (Bio-rad), through the Trans-Blot Turbo transfer system (Bio-rad). Then, the membranes were blocked in TBST (TRIS-HCl 100 mM pH 7.5, NaCl 150 mM, Tween20 0.05%) with 5% milk (Cell Signaling) to reduce nonspecific binding, washed with TBST and incubated with the primary antibodies diluted in 5% TBST BSA (Sigma Aldrich), overnight, at 4°C under slow agitation. Then, the membranes were washed and secondary antibodies (anti-rabbit XPC Cat. #MA1-23328, Thermo Scientific) were added in TBST with 5% milk (Cell Signaling), for 2 hours at room temperature and slow stirring. Subsequently, the membranes were washed for 30 minutes with TBST and the substrate Luminata (Millipore) was added for chemiluminescence detection, using the ChemiDocTM XRS equipment (Bio-rad) and analyzed with the Image Lab 3.0 program (Bio-rad). An anti-beta actin antibody (#4970S Cell Signaling) was used as a protein loading control.

### Co-Immunoprecipitation Assay

Immunoprecipitation was performed using the Pierce CO-IP Kit (Thermo Scientific) following the manufacturer’s protocol. In this experiment, 1×107 cells per IP were used. Briefly, control IgG antibodies (Cat. #2729, Cell Signaling) and mouse anti-XPC (Cat. #MA1-23328, Thermo Scientific) were immobilized using AminoLink Plus coupling resin. Equal amounts of Th17 cell lysates were pre-cleaned and subsequently incubated with the antibody-coupled resin overnight at 4°C. Subsequently, the resin was washed, and the proteins eluted with an elution buffer. Immunoprecipitated samples were analyzed for expression of BATF protein (Cat. #8638S, Cell Signaling) and XPC by immunoblotting.

### Model of colitis induced by adoptive transfer of CD4 T cells to Rag1-/- mice

CD4^+^CD45RB^high^ T cells (2.5 x 10^6^) were isolated by sorting from lymph node and spleen samples of WT and XPC^-/-^ mice, and subsequently 5 x 10^5^ cells were injected intraperitoneally into Rag1^-/-^ mice. The mice that received the cells had their weight monitored daily, and in week zero and in the fourth week after the transfer they were anesthetized and submitted to a microendoscopy procedure to evaluate inflammation in the colon. After 4 weeks, the animals were euthanized and the mesenteric lymph nodes (LNM), spleen, and colon were collected for flow cytometry and H&E analyses.

### Lamina propria cell isolation

Intestines removed from euthanized animals were opened lengthwise, washed with PBS to remove feces, and cut into pieces approximately 1 cm in size. The pieces were then incubated with RPMI medium containing 3% fetal bovine serum, EDTA, and DTT for 20 minutes at 37°C with agitation. After incubation, the pieces of intestine were sieved and washed with a medium containing EDTA to separate the intraepithelial lymphocytes. The solution resulting from the washes, which contained the intraepithelial lymphocytes, was discarded. The remaining pieces were then digested with a medium containing liberase (1/250 of a 25 mg/mL solution, Roche, Germany) and DNAse I (Sigma-Aldrich, St. Louis, MO, USA) for 25 minutes at 37°C with agitation. After digestion, the solution was passed through a sieve (70 μm), centrifuged, resuspended in PBS + 2% FBS, and passed again through a sieve (40 μm). The lamina propria cells obtained were then marked with monoclonal antibodies for analysis of the marking of surface and intracellular molecules, according to the cell population to be analyzed.

### Histological Analysis

A large intestinal fragment was removed, fixed in metacarn (60% Methanol, 30% Chloroform, and 10% Glacial Acetic Acid) for 24 hours at 4°C and subsequently transferred to a 70% alcohol solution. The material was sent for mounting in paraffin blocks and the prepared slides were stained with H&E (Hematoxylin & Eosin). Images were obtained using a Nikon microscope at 10x, 20x, and 40x. The histology score analysis was performed according to the guidance for scoring colonic inflammation mediated by disturbed immune cell homeostasis published by Erban et al. (2014).

### Alkaline comet assay

The alkaline comet assay is a single-cell electrophoresis analysis performed on days 5 after Th17 of differentiation to detect DNA breaks. The assay was conducted using the commercial kit (Trevigen, Gaithersburg, MD, USA) following the manufacturer protocol. However, the method was performed with few alterations to detect DNA oxidized bases. The DNA was digested with formamidopyrimidine DNA glycosylase (Fpg) enzyme that can identify and remove oxidized purines; this principle is often used in the comet assay detecting oxidized bases by quantifying the DNA strand breaks caused by the incubation with the enzyme (Azqueta, 2019). Briefly, 12 µl of cell suspension (1× 10 5 cells per mL) was added to 100 uL of low melting point (0.5%) agarose at 37°C and the solution was pipetted onto pre-coated CometAssay® Kit slides (Trevigen, Gaithersburg, MD, USA). After solidifying the agarose, slides were immersed in a cold lysis solution (CometAssay Lysis Solution, Trevigen, Gaithersburg, MD, USA) for at least 40 min. After lysis, slides were washed and equilibrated in an enzyme buffer (40 mM HEPES, 100 mM KCl, 0.5 mM EDTA, 0.2mg/ml BSA; pH 8). Each slide was then treated with 50 μL of enzyme buffer with or without (control) the FPG enzyme (0.12 units per gel—New England Bio Labs Inc., Ipswich, MA, EUA) and incubated in a moist box at 37 °C for 40 min. After the treatment, the slides were set at four °C for 30 min and then immersed in an alkaline solution (300 mM NaOH and one mM EDTA, pH 13.0) for 20 minutes, allowing the unfolding of the DNA structure. Later, it was submitted to an electrophoresis process in an alkaline solution (30 mM NaOH and 1 mM EDTA, pH 13.0) at 300 mA and 4°C for 30 min. Finally, slides were neutralized with dH2O for 5 min and fixed in 70% ethanol. DNA staining was done using a SYBR™ Green I Nucleic Acid Gel Stain (Invitrogen Molecular Probes, Eugene, Oregon, USA) solution for 5 min. Slides were analyzed under a Zeiss fluorescence microscope (Axiovert 200) with a 510–560 nm filter and a 590 nm barrier. DNA fragmentation parameters were analyzed using the LUCIA Comet Assay™ software (Laboratory Image, Prague, Czech Republic); 100 comets were scored from each point and the tail moment median was considered.

### Statistical analysis

The analyzes of the results were expressed as mean ± standard deviation (SD) and the following statistical tests were performed: One-way ANOVA test followed by Tukey’s post-test, or student t test followed by. Such analyzes were carried out with the help of the GraphPad Prism software (GraphPad Software Inc., San Diego CA, USA) and differences that presented p values equal to or less than 0.05 were considered statistically significant.

## Acknowledgments

We are grateful to Dr. Meire Ioshie Hiyane, for their technical assistance. We thank the Core Facility for Scientific Research at the University of São Paulo (CEFAP-USP/CONFOCAL/FLUIR/GENIAL) for the confocal microscopy, flow cytometry, and genome investigation analysis. The research leading to these results received funding from the São Paulo Research Foundation (FAPESP) under grant agreements, 2017/05264-7 and 2023/07482-2, to Niels Olsen Saraiva Câmara and 2019/19435-3 (NWO FAPESP Grant) to Carlos Menck and Niels Olsen Saraiva Câmara. Ari Waisman was supported by the Deutsche Forschungsgemeinschaft (DFG, German Research Foundation) project numbers 318346496 – SFB1292/2 and 490846870 –TRR355/1. This study was also financed in part by the São Paulo Research Foundation (FAPESP) under grant 22/13289-8 to Jefferson Antônio Leite.

## Authors contributions

J.A.L, and N.O.S.C conceived the study. J.A.L designed and performed the experiments, analyzed data and wrote the manuscript. L.M, E.M, G.S.L; N.N.B; S.B; T.G; V.Y; J.A.O. N.N; P.S; H.R, C.P.F; B.L, A.G.S; M.C helped with experiments, data analysis and discussion. T.H; S.C; L.M.C; S.M; A.W; C.F.M.M and N.O.S.C provided scientific assistance, intellectual support and critically revised the manuscript. N.O.S.C provided intellectual support in addition to directing and supervising the study. All authors have read and agreed to the published version of the manuscript.

## Data availability

The raw data from RNA-seq was deposited at Sequence read archive - SRA under the accession code: PRJNA1160718

---

**Supplementary Figure 1.**
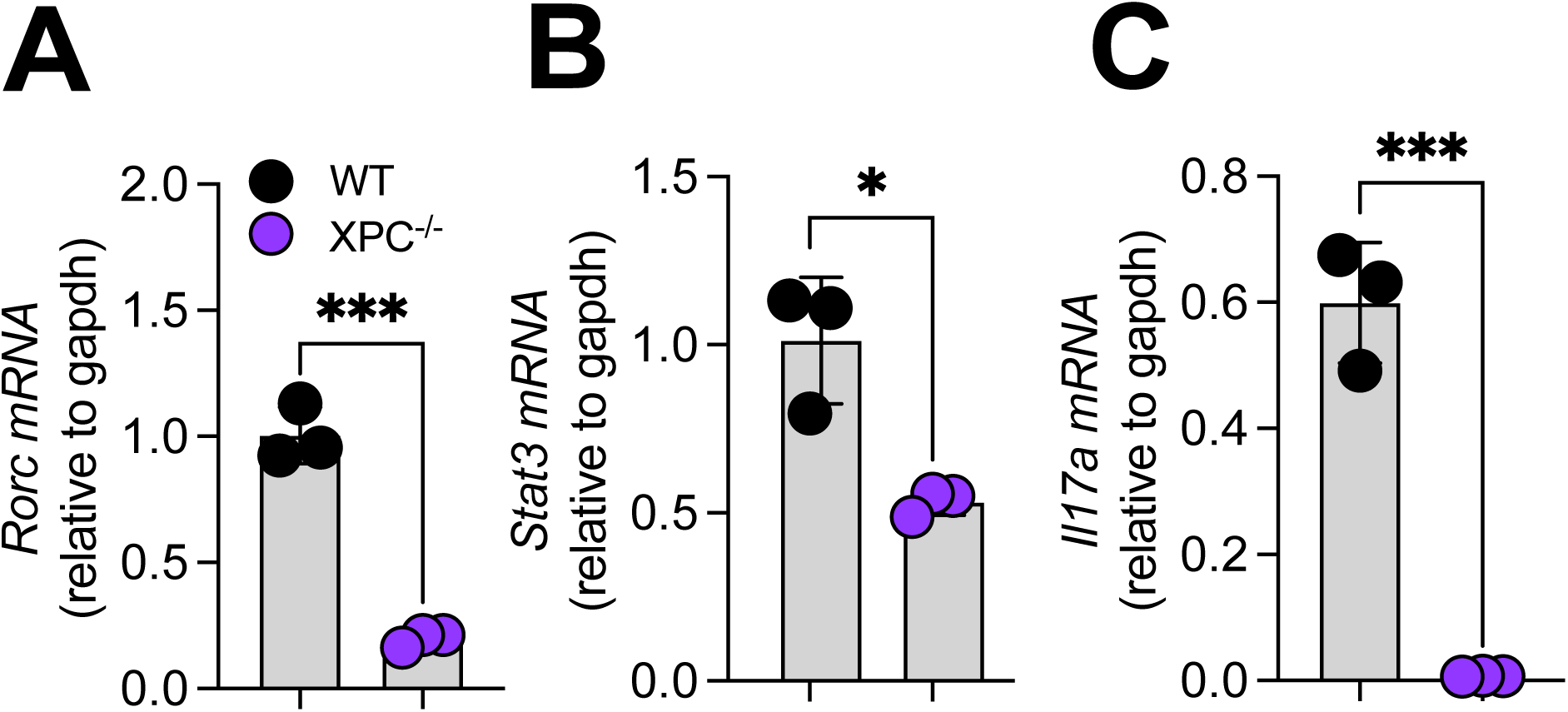
– Expression of genes associated with the Th17 program in WT and XPCKO. (A, B, C) mRNA expression of Rorc, Stat3, Il17a in WT and XPC-/- Th17 cells after 96 h of culture (n=3).

**Supplementary 2.**
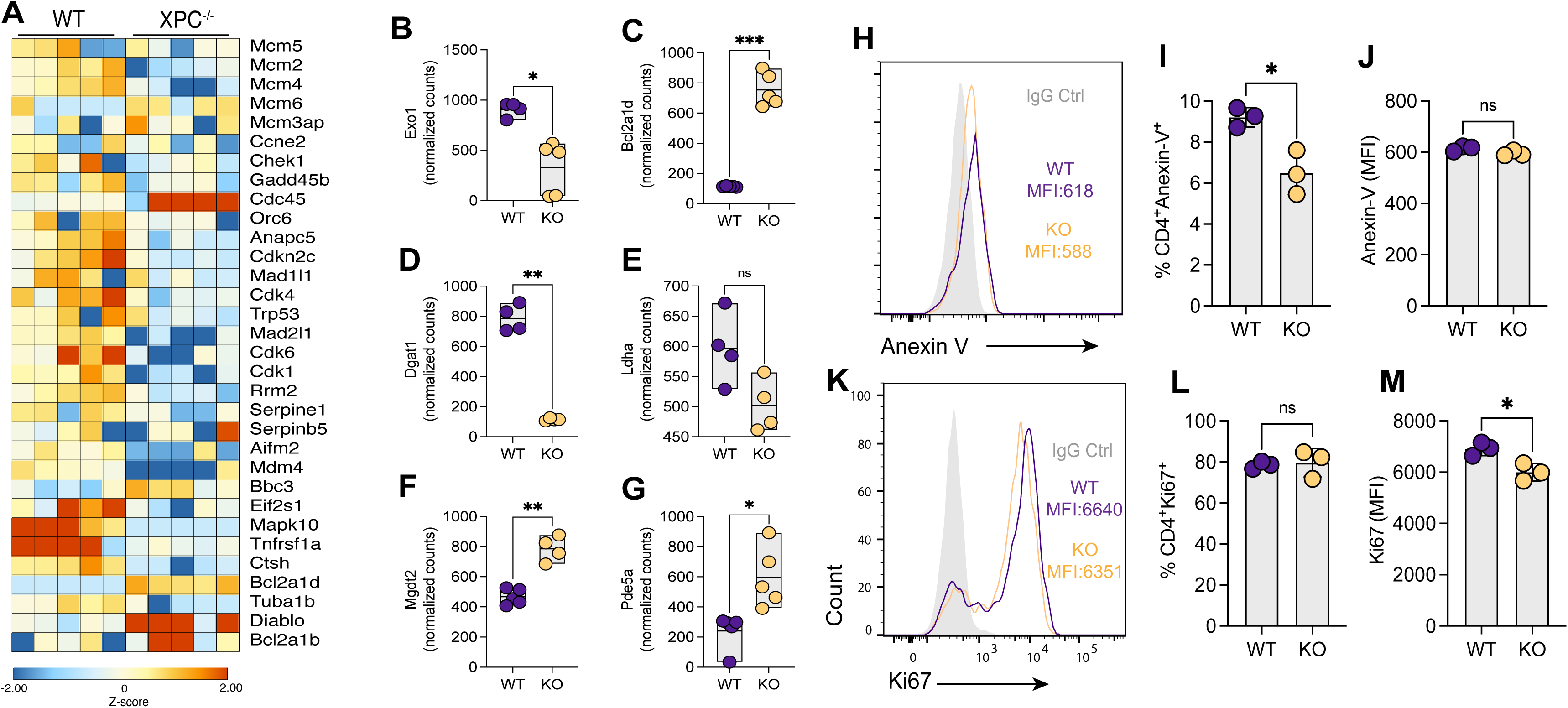
– RNA-seq analysis of WT and XPC Th17 cells and expression of Annexin V and Ki67. (A) Heat map representing the expression of genes (z-score) related to cell cycle, p53, cellular senescence and apoptosis pathways in WT and XPC^-/-^ Th17 cells; (B, C, D, E, F, G) Normalized counts of Exo1, Bcl2a1d, Dgat1, Ldha, Mgdt2 and Pde5a in WT and XPC^-/-^ Th17 cells; (H, K) Representative histogram analysis, frequency and mean of fluorescence intensity (MFI) of Annexin V in WT and XPC^-/-^ Th17 cells (n=3); (I, J, L, M) Representative histogram analysis, frequency and mean of fluorescence intensity (MFI) of Ki67 V in WT and XPC^-/-^ Th17 cells (n=3). Data are representative of at least two independent experiments and are shown as mean ± SEM. *p < 0.05 determined T-test *p < 0.05

**Supplementary Figure 3.**
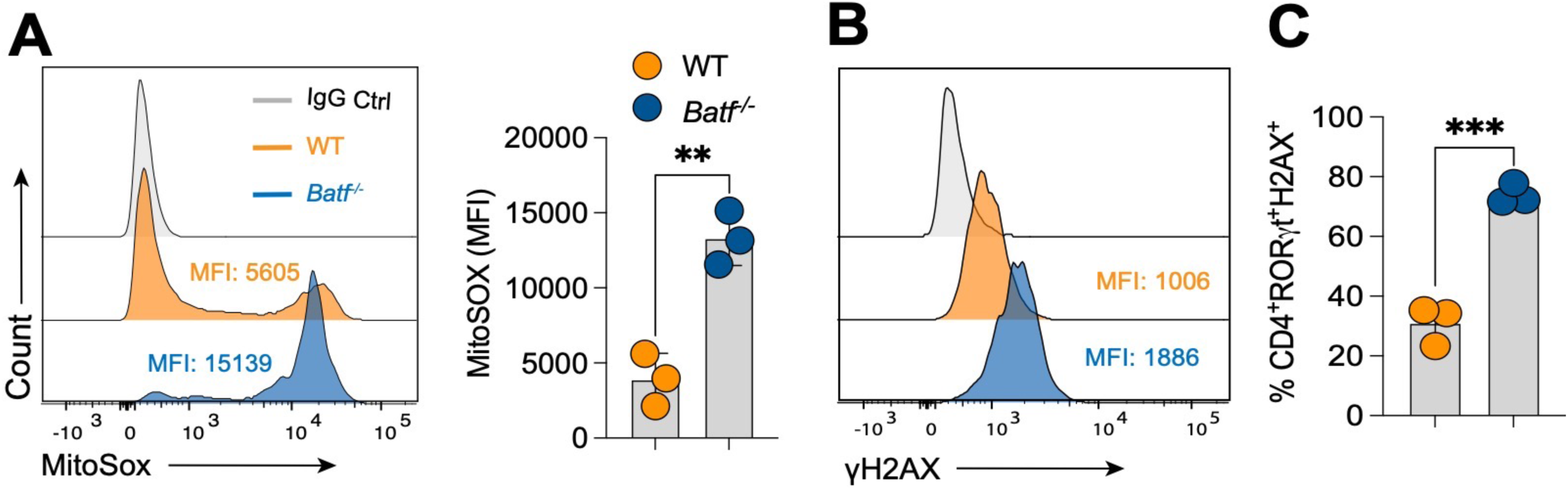
– Expression of γH2AX and MitoSOX by BATF-deficient Th17 cells. (A) Representative histogram analysis and mean of fluorescence intensity (MFI) of MitoSOX in WT and BATF^-/-^ Th17 cells (n=3); (B, C) Representative histogram analysis, frequency and mean of fluorescence intensity (MFI) of γH2AX in WT and BATF^-/-^ Th17 cells (n=3); Data are representative of at least two independent experiments and are shown as mean ± SEM. *p < 0.05 determined T-test *p < 0.05.

## References

1. McGeachy, M.J., and Cua, D.J. (2008). Th17 Cell Differentiation: The Long and Winding Road. Preprint at Immunity, 10.1016/j.immuni.2008.03.001 10.1016/j.immuni.2008.03.001.

2. Ivanov, I.I., McKenzie, B.S., Zhou, L., Tadokoro, C.E., Lepelley, A., Lafaille, J.J., Cua, D.J., and Littman, D.R. (2006). The Orphan Nuclear Receptor RORγt Directs the Differentiation Program of Proinflammatory IL-17+ T Helper Cells. Cell 126, 1121– 1133. 10.1016/J.CELL.2006.07.035/ATTACHMENT/E99CA17A-A18F-45D4-8550-15B80B8ABCF7/MMC1.PDF.

3. Veldhoen, M., Hocking, R.J., Atkins, C.J., Locksley, R.M., and Stockinger, B. (2006). TGFβ in the context of an inflammatory cytokine milieu supports de novo differentiation of IL-17-producing T cells. Immunity 24, 179–189. 10.1016/J.IMMUNI.2006.01.001/ATTACHMENT/C408AB1B-0BD6-4B45-A197-B75282853BC9/MMC1.PDF.

4. Mudter, J., Yu, J., Zufferey, C., Brüstle, A., Wirtz, S., Weigmann, B., Hoffman, A., Schenk, M., Galle, P.R., Lehr, H.A., et al. (2011). IRF4 regulates IL-17A promoter activity and controls RORγt-dependent Th17 colitis in vivo. Inflamm Bowel Dis 17, 1343–1358. 10.1002/IBD.21476.

5. Nishihara, M., Ogura, H., Ueda, N., Tsuruoka, M., Kitabayashi, C., Tsuji, F., Aono, H., Ishihara, K., Huseby, E., Betz, U.A.K., et al. (2007). IL-6-gp130-STAT3 in T cells directs the development of IL-17+ Th with a minimum effect on that of Treg in the steady state. Int Immunol 19, 695–702. 10.1093/INTIMM/DXM045.

6. Bettelli, E., Carrier, Y., Gao, W., Korn, T., Strom, T.B., Oukka, M., Weiner, H.L., and Kuchroo, V.K. (2006). Reciprocal developmental pathways for the generation of pathogenic effector TH17 and regulatory T cells. Nature 2006 441:7090 441, 235–238. 10.1038/nature04753.

7. Yang, Z., Shen, Y., Oishi, H., Matteson, E.L., Tian, L., Goronzy, J.J., and Weyand, C.M. (2016). Restoring oxidant signaling suppresses proarthritogenic T cell effector functions in rheumatoid arthritis. Sci Transl Med 8. 10.1126/SCITRANSLMED.AAD7151.

8. Li, Y., Shen, Y., Hohensinner, P., Ju, J., Wen, Z., Goodman, S.B., Zhang, H., Goronzy, J.J., and Weyand, C.M. (2016). Deficient Activity of the Nuclease MRE11A Induces T Cell Aging and Promotes Arthritogenic Effector Functions in Patients with Rheumatoid Arthritis. Immunity 45, 903–916. 10.1016/J.IMMUNI.2016.09.013.

9. Shao, L., Goronzy, J.J., and Weyand, C.M. (2010). DNA-dependent protein kinase catalytic subunit mediates T-cell loss in rheumatoid arthritis. EMBO Mol Med 2, 415–427. 10.1002/EMMM.201000096.

10. Belikov, A. V., Schraven, B., and Simeoni, L. (2015). T cells and reactive oxygen species. J Biomed Sci 22, 1–11. 10.1186/S12929-015-0194-3/FIGURES/3.

11. Zhang, J.H., Ferrante, A., Arrigo, A.P., and Dayer, J.M. (1992). Neutrophil stimulation and priming by direct contact with activated human T lymphocytes. The Journal of Immunology 148, 177–181. 10.4049/JIMMUNOL.148.1.177.

12. Shatynski, K.E., Chen, H., Kwon, J., and Williams, M.S. (2012). Decreased STAT5 phosphorylation and GATA-3 expression in NOX2-deficient T cells: role in T helper development. Eur J Immunol 42, 3202–3211. 10.1002/EJI.201242659.

13. Kaufmann, U., Kahlfuss, S., Yang, J., Ivanova, E., Koralov, S.B., and Feske, S. (2019). Calcium Signaling Controls Pathogenic Th17 Cell-Mediated Inflammation by Regulating Mitochondrial Function. Cell Metab 29, 1104–1118.e6. 10.1016/J.CMET.2019.01.019.

14. Fu, G., Xu, Q., Qiu, Y., Jin, X., Xu, T., Dong, S., Wang, J., Ke, Y., Hu, H., Cao, X., et al. (2017). Suppression of Th17 cell differentiation by misshapen/NIK-related kinase MINK1. Journal of Experimental Medicine 214, 1453–1469. 10.1084/JEM.20161120.

15. Heintzman, D.R., Elasy, J., Chi, C., Ye, X., Krystoviak, E.S., Khan, W., Olson, L., Jones, A., Voss, K., Patterson, A.R., et al. (2022). Subset-specific mitochondrial and DNA damage shapes T cell responses to fever and inflammation. bioRxiv, 2022.11.14.516478. 10.1101/2022.11.14.516478.

16. Krokan, H.E., and Bjørås, M. (2013). Base Excision Repair. Cold Spring Harb Perspect Biol 5, 1–22. 10.1101/CSHPERSPECT.A012583.

17. Mori, M.P., Costa, R.A.P., Soltys, D.T., Freire, T. de S., Rossato, F.A., Amigo, I., Kowaltowski, A.J., Vercesi, A.E., and De Souza-Pinto, N.C. (2017). Lack of XPC leads to a shift between respiratory complexes I and II but sensitizes cells to mitochondrial stress. Scientific Reports 2017 7:1 7, 1–15. 10.1038/s41598-017-00130-x.

18. D’Errico, M., Parlanti, E., Teson, M., De Jesus, B.M.B., Degan, P., Calcagnile, A., Jaruga, P., Bjørås, M., Crescenzi, M., Pedrini, A.M., et al. (2006). New functions of XPC in the protection of human skin cells from oxidative damage. EMBO Journal 25, 4305–4315. 10.1038/SJ.EMBOJ.7601277/ASSET/5F6376AA-D5C6-4CBE-8809-9ADC8AC0EE3E/ASSETS/GRAPHIC/EMBJ7601277-FIG-0009-M.JPG.

19. Fayyad, N., Kobaisi, F., Beal, D., Mahfouf, W., Ged, C., Morice-Picard, F., Fayyad-Kazan, M., Fayyad-Kazan, H., Badran, B., Rezvani, H.R., et al. (2020). Xeroderma Pigmentosum C (XPC) Mutations in Primary Fibroblasts Impair Base Excision Repair Pathway and Increase Oxidative DNA Damage. Front Genet 11, 561687. 10.3389/FGENE.2020.561687/BIBTEX.

20. Shuck, S.C., Short, E.A., and Turchi, J.J. (2008). Eukaryotic nucleotide excision repair: from understanding mechanisms to influencing biology. Cell Research 2008 18:1 18, 64–72. 10.1038/cr.2008.2.

21. Hu, Q., Xie, Y., Ge, Y., Nie, X., Tao, J., and Zhao, Y. (2018). Resting T cells are hypersensitive to DNA damage due to defective DNA repair pathway. Cell Death & Disease 2018 9:6 9, 1–11. 10.1038/s41419-018-0649-z.

22. Schnell, A., Huang, L., Singer, M., Singaraju, A., Barilla, R.M., Regan, B.M.L., Bollhagen, A., Thakore, P.I., Dionne, D., Delorey, T.M., et al. (2021). Stem-like intestinal Th17 cells give rise to pathogenic effector T cells during autoimmunity. Cell 184, 6281–6298.e23. 10.1016/J.CELL.2021.11.018.

23. Kong, L., Pokatayev, V., Lefkovith, A., Carter, G.T., Creasey, E.A., Krishna, C., Subramanian, S., Kochar, B., Ashenberg, O., Lau, H., et al. (2023). The landscape of immune dysregulation in Crohn’s disease revealed through single-cell transcriptomic profiling in the ileum and colon. Immunity 56, 444–458.e5. 10.1016/J.IMMUNI.2023.01.002.

24. Choi, E.H., and Park, S.J. (2023). TXNIP: A key protein in the cellular stress response pathway and a potential therapeutic target. Experimental & Molecular Medicine 2023 55:7 55, 1348–1356. 10.1038/s12276-023-01019-8.

25. Manolakou, T., Verginis, P., and Boumpas, D.T. (2021). DNA Damage Response in the Adaptive Arm of the Immune System: Implications for Autoimmunity. International Journal of Molecular Sciences 2021, Vol. 22, Page 5842 22, 5842. 10.3390/IJMS22115842.

26. McNally, J.P., Millen, S.H., Chaturvedi, V., Lakes, N., Terrell, C.E., Elfers, E.E., Carroll, K.R., Hogan, S.P., Andreassen, P.R., Kanter, J., et al. (2017). Manipulating DNA damage-response signaling for the treatment of immune-mediated diseases. Proc Natl Acad Sci U S A 114, E4782–E4791. 10.1073/PNAS.1703683114.

27. Alissafi, T., Kalafati, L., Lazari, M., Filia, A., Kloukina, I., Manifava, M., Lim, J.H., Alexaki, V.I., Ktistakis, N.T., Doskas, T., et al. (2020). Mitochondrial Oxidative Damage Underlies Regulatory T Cell Defects in Autoimmunity. Cell Metab 32, 591–604.e7. 10.1016/J.CMET.2020.07.001.

28. Nasrallah, N. Al, Wiese, B.M., and Sears, C.R. (2022). Xeroderma Pigmentosum Complementation Group C (XPC): Emerging Roles in Non-Dermatologic Malignancies. Front Oncol 12. 10.3389/FONC.2022.846965.

29. Bradford, P.T., Goldstein, A.M., Tamura, D., Khan, S.G., Ueda, T., Boyle, J., Oh, K.S., Imoto, K., Inui, H., Moriwaki, S.I., et al. (2011). Cancer and neurologic degeneration in xeroderma pigmentosum: long term follow-up characterises the role of DNA repair. J Med Genet 48, 168–176. 10.1136/JMG.2010.083022.

30. Mariani, E., Facchini, A., Honorati, M.C., Lalli, E., Berardesca, E., Ghetti, P., Marinoni, S., Nuzzo, F., Ricotti, G.C.B.A., and Stefanini, M. (1992). Immune defects in families and patients with xeroderma pigmentosum and trichothiodystrophy. Clin Exp Immunol 88, 376. 10.1111/J.1365-2249.1992.TB06457.X.

31. Abid, K., El Mezni, F., Kamoun, M.R., Fazaa, B., Zermani, R., Hadouchi, C., and Hamzaoui, K. (2010). Xeroderma pigmentosum skin: an immune privilege site for tumor development. J Cutan Pathol 37, 452–459. 10.1111/J.1600-0560.2009.01401.X.

32. Lai, J.P., Liu, Y.C., Alimchandani, M., Liu, Q., Aung, P.P., Matsuda, K., Lee, C.C.R., Tsokos, M., Hewitt, S., Rushing, E.J., et al. (2014). The influence of DNA repair on neurological degeneration, cachexia, skin cancer and internal neoplasms: Autopsy report of four xeroderma pigmentosum patients (XP-A, XP-C and XP-D). Acta Neuropathol Commun 2, 1–17. 10.1186/2051-5960-1-4/FIGURES/9.

33. Ren, Y., Ma, J., Wang, T., Bu, R., Kong, X., Shi, Y., and Zhang, L. (2018). Interleukin-17 inhibits the growth of oral squamous cell carcinoma by promoting the differentiation of T helper 17 cells. Transl Cancer Res 7, 839–848. 10.21037/TCR.2018.06.05.

34. Muranski, P., Boni, A., Antony, P.A., Cassard, L., Irvine, K.R., Kaiser, A., Paulos, C.M., Palmer, D.C., Touloukian, C.E., Ptak, K., et al. (2008). Tumor-specific Th17-polarized cells eradicate large established melanoma. Blood 112, 362–373. 10.1182/BLOOD-2007-11-120998.

35. von Euw, E., Chodon, T., Attar, N., Jalil, J., Koya, R.C., Comin-Anduix, B., and Ribas, A. (2009). CTLA4 blockade increases Th17 cells in patients with metastatic melanoma. J Transl Med 7, 1–13. 10.1186/1479-5876-7-35/FIGURES/4.

36. Iwakura, Y., Nakae, S., Saijo, S., and Ishigame, H. (2008). The roles of IL-17A in inflammatory immune responses and host defense against pathogens. Immunol Rev 226, 57–79. 10.1111/J.1600-065X.2008.00699.X.

37. Sugasawa, K., Akagi, J. ichi, Nishi, R., Iwai, S., and Hanaoka, F. (2009). Two-step recognition of DNA damage for mammalian nucleotide excision repair: Directional binding of the XPC complex and DNA strand scanning. Mol Cell 36, 642–653. 10.1016/J.MOLCEL.2009.09.035.

38. Shell, S.M., Hawkins, E.K., Tsai, M.S., Hlaing, A.S., Rizzo, C.J., and Chazin, W.J. (2013). Xeroderma pigmentosum complementation group C protein (XPC) serves as a general sensor of damaged DNA. DNA Repair (Amst) 12, 947–953. 10.1016/J.DNAREP.2013.08.013.

39. Mu, H., Geacintov, N.E., Broyde, S., Yeo, J.E., and Schärer, O.D. (2018). Molecular basis for damage recognition and verification by XPC-RAD23B and TFIIH in nucleotide excision repair. DNA Repair (Amst) 71, 33–42. 10.1016/J.DNAREP.2018.08.005.

40. Cobley, J.N., Margaritelis, N. V., Morton, J.P., Close, G.L., Nikolaidis, M.G., and Malone, J.K. (2015). The basic chemistry of exercise-induced DNA oxidation: Oxidative damage, redox signaling, and their interplay. Front Physiol 6. 10.3389/FPHYS.2015.00182.

41. Fleming, A.M., and Burrows, C.J. (2022). Chemistry of ROS-mediated oxidation to the guanine base in DNA and its biological consequences. Int J Radiat Biol 98, 452–460. 10.1080/09553002.2021.2003464.

42. Charlet-Berguerand, N., Feuerhahn, S., Kong, S.E., Ziserman, H., Conaway, J.W., Conaway, R., and Egly, J.M. (2006). RNA polymerase II bypass of oxidative DNA damage is regulated by transcription elongation factors. EMBO Journal 25, 5481–5491. 10.1038/SJ.EMBOJ.7601403/ASSET/08D41866-AAF1-4562-A687-D4E8AAD30C8C/ASSETS/GRAPHIC/EMBJ7601403-FIG-0004-M.JPG.

43. de Melo, J.T.A., de Souza Timoteo, A.R., Lajus, T.B.P., Brandão, J.A., de Souza-Pinto, N.C., Menck, C.F.M., Campalans, A., Radicella, J.P., Vessoni, A.T., Muotri, A.R., et al. (2016). XPC deficiency is related to APE1 and OGG1 expression and function. Mutat Res 784–785, 25–33. 10.1016/J.MRFMMM.2016.01.004.

44. Bidon, B., Iltis, I., Semer, M., Nagy, Z., Larnicol, A., Cribier, A., Benkirane, M., Coin, F., Egly, J.M., and Le May, N. (2018). XPC is an RNA polymerase II cofactor recruiting ATAC to promoters by interacting with E2F1. Nat Commun 9. 10.1038/S41467-018-05010-0.

45. Denechaud, P.D., Fajas, L., and Giralt, A. (2017). E2F1, a novel regulator of metabolism. Front Endocrinol (Lausanne) 8, 307580. 10.3389/FENDO.2017.00311/BIBTEX.

46. Graves, J.D., Lee, Y.J., Liu, K., Li, G., Lin, F.T., and Lin, W.C. (2020). E2F1 sumoylation as a protective cellular mechanism in oxidative stress response. Proc Natl Acad Sci U S A 117, 14958–14969. 10.1073/PNAS.1921554117/SUPPL_FILE/PNAS.1921554117.SAPP.PDF.

47. Le May, N., Mota-Fernandes, D., Vélez-Cruz, R., Iltis, I., Biard, D., and Egly, J.M. (2010). NER factors are recruited to active promoters and facilitate chromatin modification for transcription in the absence of exogenous genotoxic attack. Mol Cell 38, 54–66. 10.1016/J.MOLCEL.2010.03.004.

48. Schraml, B.U., Hildner, K., Ise, W., Lee, W.L., Smith, W.A.E., Solomon, B., Sahota, G., Sim, J., Mukasa, R., Cemerski, S., et al. (2009). The AP-1 transcription factor Batf controls TH17 differentiation. Nature 460, 405. 10.1038/NATURE08114.

49. Pham, D., Silberger, D.J., Nguyen, K.N., Gao, M., Weaver, C.T., and Hatton, R.D. (2023). Batf stabilizes Th17 cell development via impaired Stat5 recruitment of Ets1-Runx1 complexes. EMBO J 42. 10.15252/EMBJ.2021109803.

